# Uncovering High-Resolution Organization of Genomic Loci using Experimentally Informed Polymer Model

**DOI:** 10.1101/2025.11.19.689404

**Authors:** Rahul Mittal, Dieter W. Heermann, Arnab Bhattacherjee

## Abstract

The functionality of an organism depends on gene expression patterns, which are governed by the hierarchical organization of the genome. Numerous efforts, leveraging both polymer physics-based models and experimental imaging technologies, have sought to elucidate the structure-function relationship of chromatin fibers. However, a major challenge remains: the intrinsically multi-scale nature of chromatin organization, which has yet to be fully explored. Here, we present an experimentally informed, polymer physics-based model capable of reconstructing chromatin structural ensembles by integrating low-resolution contact data with MNase-derived nucleosome positioning information. We applied this model to multiple human genomic loci. Our analysis uncovered distinct structural features associated with active and inactive chromatin states, providing key insights into the relationship between genomic organization and transcriptional activity. These findings offer a promising framework for understanding genome structure-function relationships and their implications in developmental disorders and gene misregulation-related diseases.

## 1 Introduction

Accurate decoding of genetic information from DNA sequences is fundamental for executing essential cellular programs and responding effectively to cellular challenges [1–3]. This process initiates with the identification of specific DNA sequences by DNA-binding proteins (DBPs), followed by their precise binding. While a confluence of factors, encompassing local DNA geometry [4–6], sequence [7, 8], structure [9–11], and the concentration of the searching and other proteins [12–14] regulate the target search process of DBPs, the search efficiency primarily hinges on the accessibility of the cognate DNA sites to the DBPs [15]. This accessibility is particularly critical in eukaryotes, where the entire genome must be tightly and hierarchically packaged within the confines of the cell nucleus [16–20].

At the primary level, genomic DNA is wrapped around an octamer of histone proteins, forming nucleosomes, where approximately 147 base pairs of DNA are tightly wound into superhelical turns [18, 21]. This association between nucleosomal DNA and the histone core significantly restricts the access of DNA sites to non-histone proteins, thus serving as a crucial regulatory layer in determining cell identity and controlling gene expression [22]. Importantly, nucleosomes are not static structures; their dynamic behavior, including breathing [14, 23] and sliding [24, 25], provides transient access to DBPs to look for their cognate DNA sites. Furthermore, nucleosome positioning along the chromatin fiber can exhibit distinct patterns [26–28]: phased arrays with regularly spaced nucleosomes, unphased arrays with less regular spacing, and fuzzy arrays with more irregular positioning. These spatial arrangements contribute to the folding of chromatin into higher-order structures, which are essential for its overall functionality. However, the fundamental principles governing this organization remain unclear, largely due to the vast range of length scales involved. This complexity poses a significant challenge in assessing how higher-order chromatin architecture influences gene function.

The advent of next-generation sequencing (NGS) technologies, particularly, the chromosome conformation capture (3C) family of methods [29–31] such as Hi-C [32], have provided critical insights regarding chromatin organization [33–35] at a kbp to Mbp length scale. These studies based on pairwise contact frequencies between the segments of genome have revealed the existence of topologically associated domains (TADs) [36–38] and chromatin loops [39–41]. The abundance of TADs across the genome of different species confirms that they are a conserved across the genome and serve as architectural units of the chromatin fibre that define regulatory landscapes and play a crucial role in shaping functional chromosomal organization. TAD boundaries represent the replication domains [42], and genes within the domain are coregulated during cell differentiation [43, 44]. The significance of TAD is further manifested from the fact that disruption of TAD structures due to modulation in their boundaries can lead to ectopic contacts between cis-regulating elements and gene promoters. This leads to misexpression of genes, which is closely related to developmental defects and cancer [45, 46]. The major drawback of these methods lies in their inability to provide detailed structural information about chromatin organization. To address this, various theoretical approaches have been developed to complement experimental findings and explore the structure-function relationship of chromatin. These methods predominantly fall into two categories: data-driven models [47–52] and polymer physics-based simulations [53–60]. Majority of the approaches utilize experimental Hi-C contact maps as input, employing multi-parameter polymer models to iteratively optimize interactions between genomic loci to reproduce experimental contact frequencies. Other polymer physics-based models simulate chromatin organization using principles such as cohesin-mediated [61] loop extrusion [41, 62, 63], protein-bridging interactions [55, 56], or phase separation driven by specific epigenetic modifications [52, 59]. While these models provide valuable insights, they typically resolve chromatin organization at megabase (Mbp) scales due to the coarse-grained resolution of input Hi-C data excluding the possibility of resolving large-scale organization and probe its influence on the fine-grained regulatory mechanisms driving gene expression.

To overcome this limitation, we present an experimentally informed polymer model that integrates Hi-C contact data with nucleosome positioning information to predict the large-scale architecture of the chromatin fiber at near base-pair-level resolution. The model employs a two-tiered approach to bridge the disparate length scales of chromatin organization. At the first level, the model generates an ensemble of coarse-grained chromatin conformations using Hi-C data as input, capturing large-scale architectural features. These coarse conformations then serve as structural frameworks for the second level, where high-resolution polymer conformations are modeled. The fine-scale resolution incorporates nucleosomes and linker DNA segments, guided by nucleosome positioning data derived from MNase-seq experiments. Applying this method to 0.2 Mbp genomic stretches encompassing the Nanog and HoxB4 loci in human stem cells (hSCs), we find that nucleosome condensates constitute a critical organizational feature of the chromatin fiber. The morphology of these condensates, which we refer to as “nucleosome blobs,” closely resembles the blob-like structures observed in live super-resolution imaging of human bone osteosarcoma (U2OS) cells [64]. Moreover, our results reveal that at the local scale, the internal organization of nucleosomes within these blobs differs markedly between the Nanog and HoxB4 loci. At a global scale, the spatial distribution of nucleosome blobs around these loci exhibits distinct patterns that influence both the free energy landscape of the genomic regions and the physiological properties of the chromatin fiber. These findings offer a compelling physical framework for understanding the transcriptional activity at the Nanog and HoxB4 loci, paving the way for deeper insights into the structure-function relationships of chromatin fibers. This method may prove instrumental in unraveling the causal regulatory mechanisms governing gene expression, development, and disease pathogenesis.

## 2 Results

### 2.1 Experimentally informed polymer model of chromatin

To investigate large-scale chromatin organization and its impact on the intricate regulatory mechanisms governing gene expression, we used a multi-step method to generate high-resolution chromatin conformations. In the first stage, we generate an ensemble of steady-state chromatin configurations for a 0.2 Mbps segment of chromosome VII of human embryonic stem cells at a resolution of 5 kbps by considering the contact probabilities (*P*_*ij*_) of an experimental Hi-C map as input (Fig. 1A, Methods, and Supplementary Information). This approach draws upon a method proposed by Kadam et al. [65], which supports scale-dependent systematic coarse-grained simulations of chromatin fibers. The ensemble of chromatin configurations generated in this stage serve as conformational constraints for the second stage of generating fine-grained chromatin conformations. It should be noted that the first step is crucial, as the choice of input contact map plays a pivotal role in accurately capturing large-scale chromatin organization by implicitly reflecting the influence of architectural proteins.

**Fig. 1.**
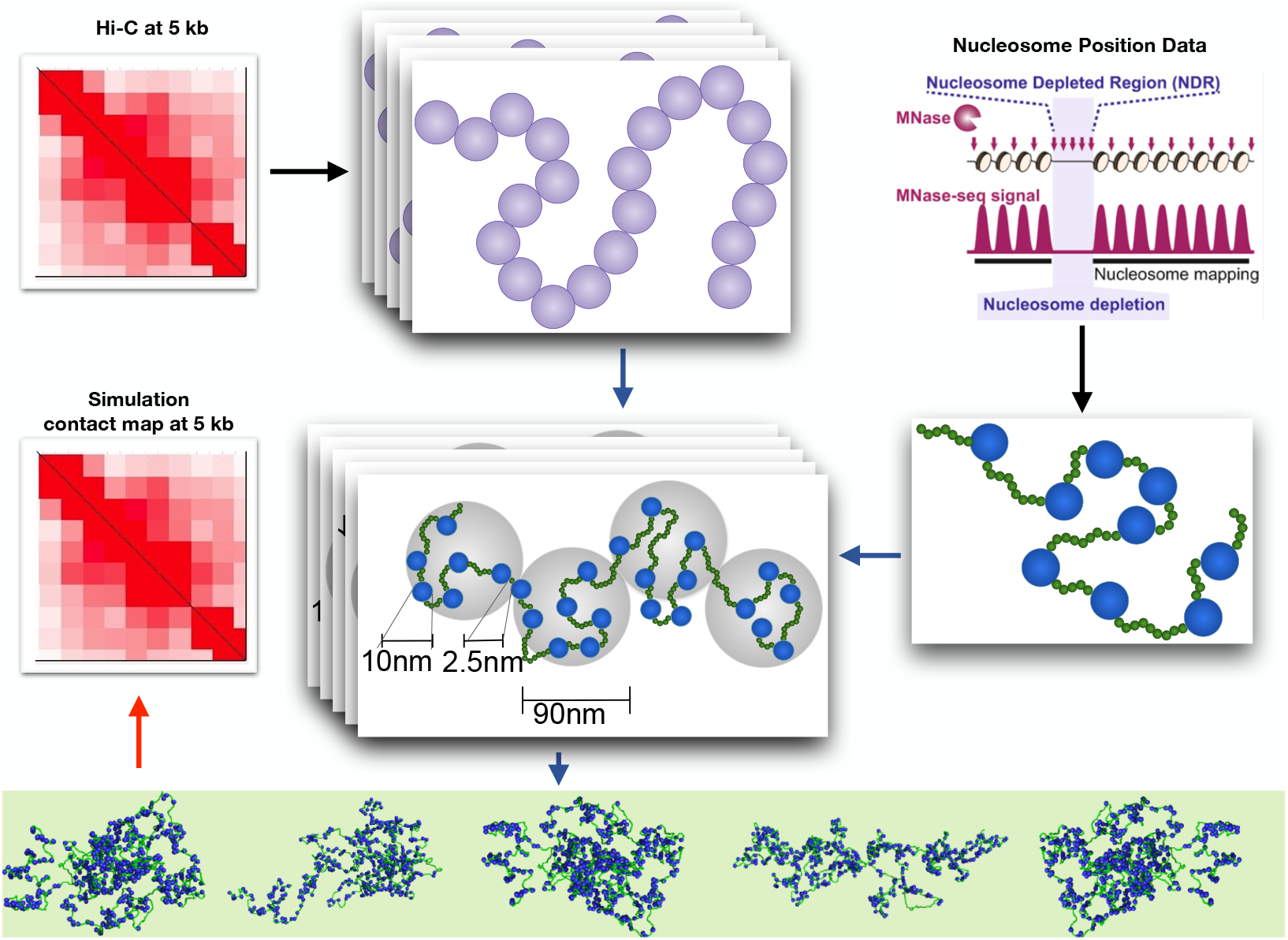
Schematic representation of the experimentally informed polymer model of chromatin. The method consists of two key stages. In the first stage, a Hi-C contact map at 5 kb resolution is used to generate multiple chromatin conformations consistent with Hi-C data. In the second stage, the ensemble of chromatin layouts is integrated with MNase-Seq data, which defines nucleosome positions, to generate NL-resolution chromatin conformations. To validate the model, ensemble-averaged contact maps at the desired resolutions are computed and compared against experimental Hi-C and Micro-C contact maps.

For the fine-grained representation of the chromatin segment, we adopt the “nucleosome-linker” (NL) bead model. This approach uses data from micrococcal nuclease digestion followed by sequencing (MNase-seq) [66] to infer nucleosome positions and linker lengths (Fig. 1B, Methods and Supplementary Information). Although nucleosome positions naturally exhibit cell-to-cell variability, we simplify our model by considering a single set of “most likely” nucleosome positions, determined using the DANPOS [67, 68] software. The NL-bead polymer chain was then simulated to fold according to the coarse-grained conformational constraints derived in the first stage. Following this initial folding, each NL-bead polymer chain undergoes further simulations to produce a large ensemble of high-resolution chromatin conformations. The adoption of the NL-bead description for chromatin fiber is motivated by insights from two key studies. The first, by Weise et al. [69], demonstrates that nucleosome positioning data is sufficient to accurately predict patterns of chromatin interactions and domain boundaries observed in experimental studies. Moreover, it highlights that nucleosome spacing significantly influences the larger-scale domain structure of chromatin. The second study, conducted by our group, confirms that the patterns in nucleosome positioning data extend beyond the conventional distinctions of heterochromatin and euchromatin, uncovering additional chromatin states that regulate differential gene expression [27]. Furthermore, conserved distribution patterns of nucleosomes along the chromatin fiber, both across the genome and within species, suggest that nucleosomes play a critical role in hierarchical chromatin organization.

### 2.2 The model precisely captures the architectural details of chromatin organization

Another significant advantage of adopting the NL-bead-level description for the chromatin fiber is its adaptability, enabling coarse-graining to match the resolutions of Micro-C and Hi-C experimental contact maps. This flexibility allows for effective validation of our model across different length scales. To this end, we analyzed two additional chromatin segments: a 0.2 Mbps region encompassing the Nanog locus on chromosome XII and a 0.2 Mbps region near the HoxB4 locus on chromosome XVII. These were tested along with the 0.2 Mbps segment of chromosome VII of human embryonic stem cells, offering a robust evaluation of the generality of our model. For all three chromatin segments, an ensemble of 13000 structures was generated at the NL-bead-level description, with representative snapshots of the systems shown in Fig. 4A-C.

We coarse-grained these conformations at resolutions of 200 bps and 5 kbp, respectively, and constructed contact maps (see Supplementary information). Ensemble-averaged contact maps were produced by averaging the individual maps from the generated conformations at both resolutions and are presented alongside available experimental Micro-C and Hi-C contact maps in Fig. 2A-F. To quantify the agreement between the simulation-generated ensemble-averaged contact maps and the experimental data, we calculated both Pearson and Spearman correlation coefficients. Notably, using both metrics provides a more comprehensive understanding of the relationship between two contact maps: Pearson captures linear trends, while Spearman identifies monotonic relationships and is robust to outliers and nonnormal data. This dual approach ensures the results are not biased by the limitations of a single method. As shown in Fig. 2A-F and in Table 1, our results demonstrate that, for all three chromatin segments, the model successfully generated ensembles of conformations that accurately reproduce the ensemble-averaged experimental contact maps across different length scales.

**Table 1.** Comparison of chromatin structural agreement metrics at different resolutions (5 kbps and 200 bps) for three chromosome segments (chrVII, chrXII, and chrXVII). The table presents Kullback-Leibler (KL) divergence, Spearman correlation, and Pearson correlation values, which assess the similarity between simulated and experimental contact maps. Lower KL divergence and higher correlation values indicate better agreement with experimental data.

| Project | 5kbps |  |  | 200bps |  |  |
| --- | --- | --- | --- | --- | --- | --- |
|  | chrVII | chrXII | chrXVII | chrVII | chrXII | chrXVII |
| KL divergence | 0.047 | 0.055 | 0.057 | 0.265 | 0.312 | 0.221 |
| Spearman Correlation | 0.911 | 0.851 | 0.927 | 0.911 | 0.931 | 0.876 |
| Pearson Correlation | 0.979 | 0.971 | 0.978 | 0.802 | 0.774 | 0.845 |

**Fig. 2.**
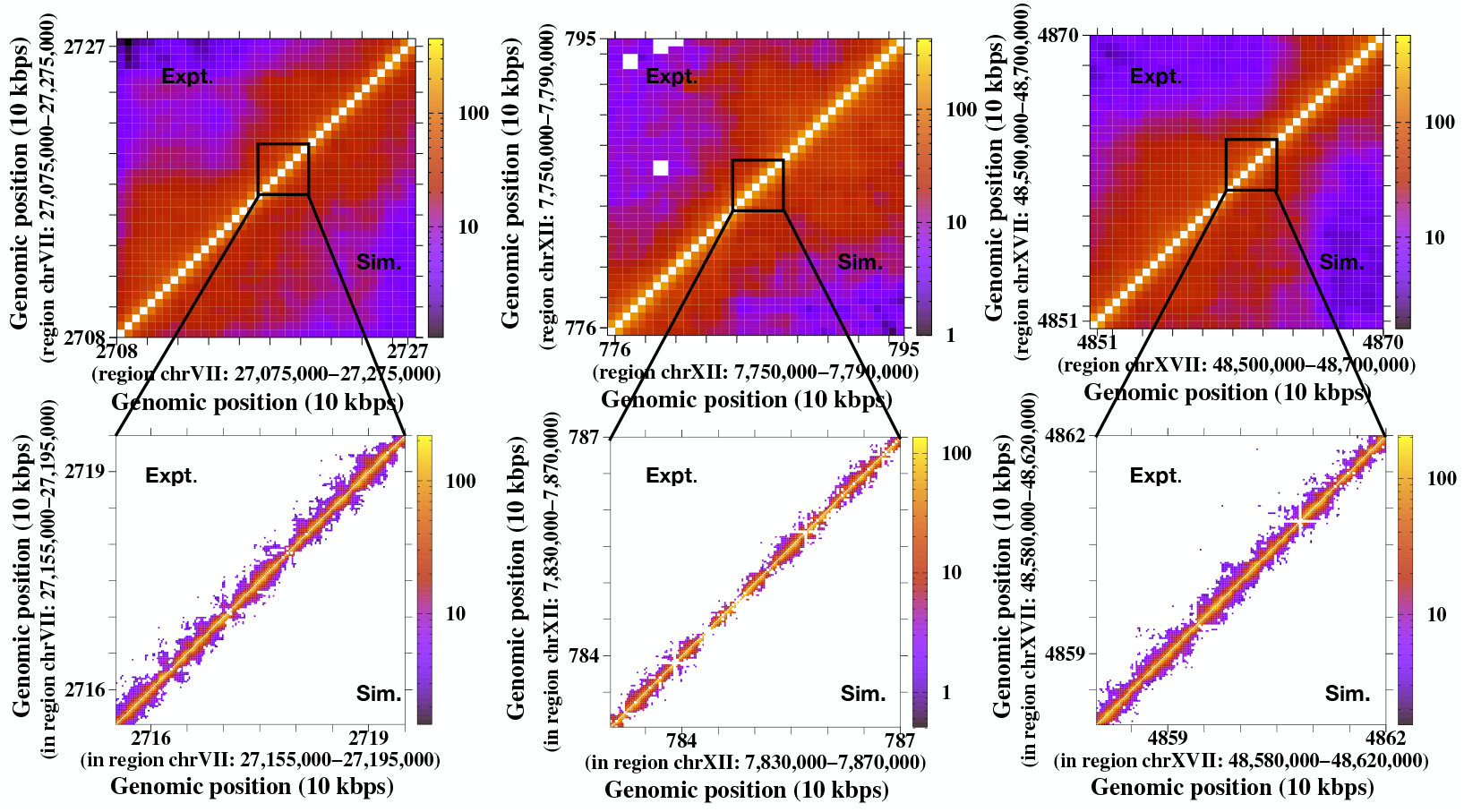
Comparison between experimental and simulated contact maps for different chromatin segments of hESCs. (A–C) show the agreement between Hi-C and simulated contact maps at 5 kb resolution for 0.2 Mb chromatin segments from human chromosomes VII, XII, and XVII, respectively. (D–F) highlight the similarities between Micro-C and simulated contact maps at 200 bp resolution for randomly selected 30 kb regions within the same chromatin segments.

We further estimated the contact frequency at 200 bps resolution as a function of genomic separation and presented them in Fig. 3A-C along with their experimental results. Our results suggest an excellent agreement with the experimental contact frequency, where short-ranged interactions are prevalent compared to less frequent long-ranged contacts.

**Fig. 3.**
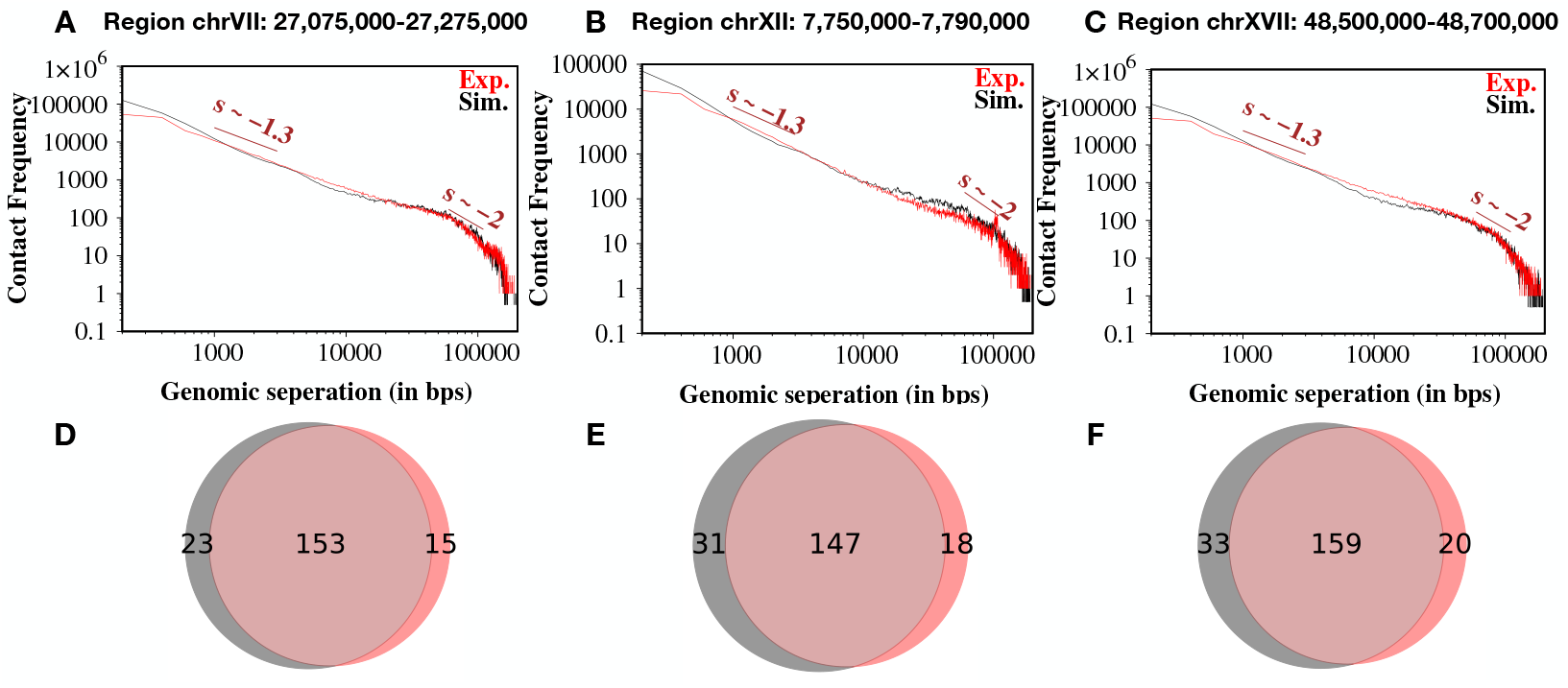
Micro-C contact frequencies as a function of genomic separation and prediction of domain boundaries. (A–C) depict the agreement between variations in Micro-C and simulated contact frequencies at 200 bp resolution as a function of genomic separation for three chromatin segments of hESCs. The measured slopes reveal distinct patterns in short-to-intermediate-range and long-range contacts within the Micro-C data. (D–F) show Venn diagrams comparing the predicted domain boundaries identified in Micro-C (red) and simulated contact maps (gray) at 200 bp resolution, highlighting their overlap and differences.

Another intriguing feature observed in the representative snapshots (Fig. 4A-C) of the three chromatin segments is the presence of irregularly spaced nucleosomes forming spatially heterogeneous blobs. This arrangement is reminiscent of the nucleosome “clutches” reported in imaging experiments on human cells [70]. On Micro-C contact maps, these blobs appear as domains characterized by densely connected squares. The separation between two squares represents the boundary between domains. To further validate our simulation results against experimental data, we identified domains by detecting boundaries (see Supplementary Text). In the 0.2 Mbps segment of human chromosome VII, the Micro-C data reveals 121 boundaries, of which our simulation-generated ensemble-averaged contact map accurately predicts 112 boundaries (see Fig. 3D), achieving a remarkable success rate of 92%. Additionally, the simulations predict 15 boundaries that are not observed experimentally. Extending this analysis to all three simulated regions, our simulations demonstrate an impressive achievement (see Fig. 3D-F), correctly predicting roughly 90% of the total boundaries (459 out of 512).

**Fig. 4.**
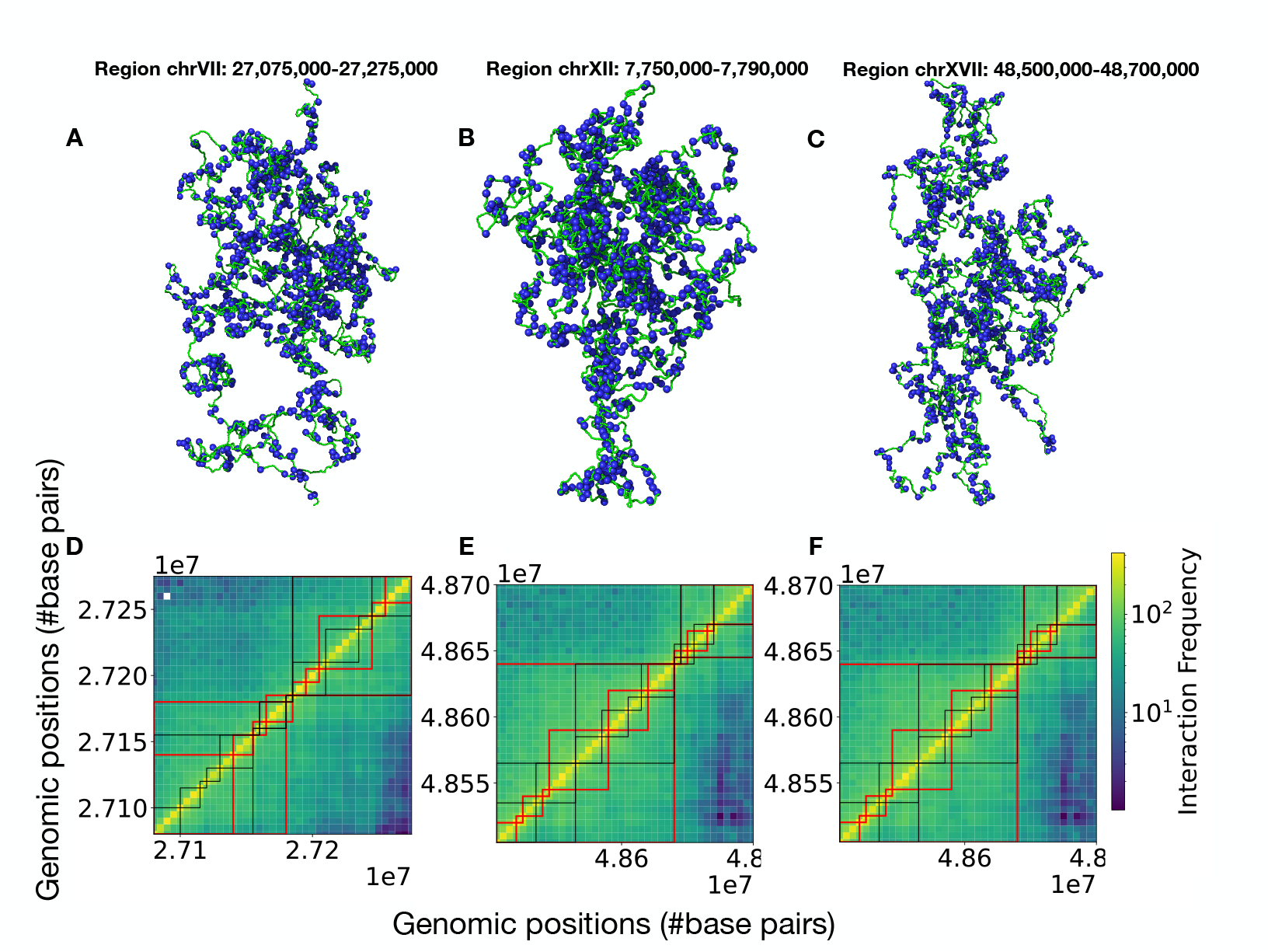
Identification of nucleosome blobs and their role in shaping the contact frequency landscape. (A–C) illustrate representative conformations of three distinct chromatin segments in hESCs, highlighting the formation of nucleosome clusters as blobs. These nucleosome blobs are identified using the DBSCAN (Density-Based Spatial Clustering of Applications with Noise) method, which distinguishes regions of high nucleosome density from areas of low nucleosome density. (D–F) demonstrate that intra-blob connectivity effectively captures the sub-TAD organization observed in experimental Hi-C maps, emphasizing the pivotal role of nucleosome blobs in shaping chromatin conformational features.

### 2.3 Nucleosome blobs are nonrandom critical component of chromatin architecture

Despite the simplistic NL-bead representation of the chromatin segments—where nucleosomes are modeled as spheres rather than their more realistic disk-like shape, and complex inter-nucleosome interactions mediated by histone tails are disregarded—our simulations effectively capture essential details of both short- and long-range contacts as shown in Fig. 2 and Fig. 3 of the previous section. The validation emphasizes its potential for uncovering deeper insights into the organizational architecture of chromatin segments.

To advance this exploration, we selected 0.2 Mbps segments of the Nanog and HoxB4 genomic loci on chromosomes XII and XVII, respectively, in human stem cells (hSCs). The rationale for this selection lies in the markedly different transcriptional activities of these two loci within this particular cell line, as evidenced by ChromHMM analysis (see Supplementary text). ChromHMM annotates genomic regions with distinct chromatin states by integrating epigenetic information and chromatin datasets using a multivariate Hidden Markov Model. For this study, we utilized the Roadmap Epigenomics Expanded Model to identify chromatin states in our target regions. This model categorizes chromatin into 18 distinct states [71–73], where states 1–12 are considered active, and states 13–18 are considered inactive [66–68, 74, 75]. The classification clearly demonstrates that Nanog, a homeobox transcription factor, functions as a transcriptionally active genomic locus. It plays a pivotal role in maintaining embryonic stem cells (ESCs) and contributes significantly to cancer development. In contrast, although HoxB4 exhibits regulatory roles in stem cells, it remains transcriptionally inactive in this specific hSC line.

To probe the organizational architecture of these two genomic loci, we analyzed an ensemble of 13000 snapshots generated from our NL-bead simulations. As shown in the respective snapshots in Fig. 4A-C, we begin by investigating the clustering of nucleosomes in the form of blobs. This is done with the DBSCAN (Density-Based Spatial Clustering of Applications with Noise) method [76] that detects nucleosome blobs. This powerful clustering algorithm effectively identifies clusters (or blobs) as regions of high nucleosome density separated by areas of low nucleosome density. DBSCAN is particularly adept at detecting blobs with irregular or non-convex shapes and differentiates between noise (low-density points) and significant nucleosome clusters, providing a nuanced understanding of the spatial organization. We applied this method to identify blobs across 13,000 chromatin conformations, where each blob consists of multiple closely interacting nucleosomes. To assess the significance of these interactions, we computed the contact frequency for each unique nucleosome pair within the same blob for blobs containing 50 or more nucleosomes. The resulting contact frequency map was then compared with the Hi-C contact map, as shown in Fig. 4D–F for all three selected chromatin segments. The results reveal a strong correlation (roughly than 90%) between Hi-C contacts and intra-blob nucleosome interactions for all the three chromatin segments, suggesting that blobs are not random assemblies but rather fundamental architectural units of chromatin organization. Furthermore, we analyzed the boundaries in these contact maps and found that while the intra-blob contact information accurately captures the domain-like organizations in the experimental contact map (indicated by red lines), it also reveals sub-TAD organizations (indicated by black lines) that would otherwise remain undetectable at a 5 kbps Hi-C resolution.

### 2.4 Physiology of nucleosome blobs in Nanog and HoxB4 genomic loci

Next, we proceed to characterize the physiology of the nucleosome blobs for both Nanog and HoxB4 genomic loci. The distributions of surface area of blobs presented in Fig. 5A, measured using Convex Hull algorithms [77], fit a log-normal distribution for both the genomic loci with parameters 6.9 ×10^−3^ ± 0.019*µm*^2^ for Nanog blobs and 7.1 × 10^−3^ ± 0.018*µm*^2^ for Nanog blobs. Their size distribution in terms of base pairs (see Fig S5) indicates the most probable blobs arises due to accumulation of 660 bps that includes multiple nucleosomes. It is noteworthy that the estimation of surface area of the blobs is in excellent agreement with the surface area observed for blobs in chromatin for live and fixed human cells observed through high-density photoactivated localization microscopy [64, 78]. The high value of standard deviation in the surface area distributions indicates large variability in blob sizes with a long tail in the area distribution toward large values.

**Fig. 5.**
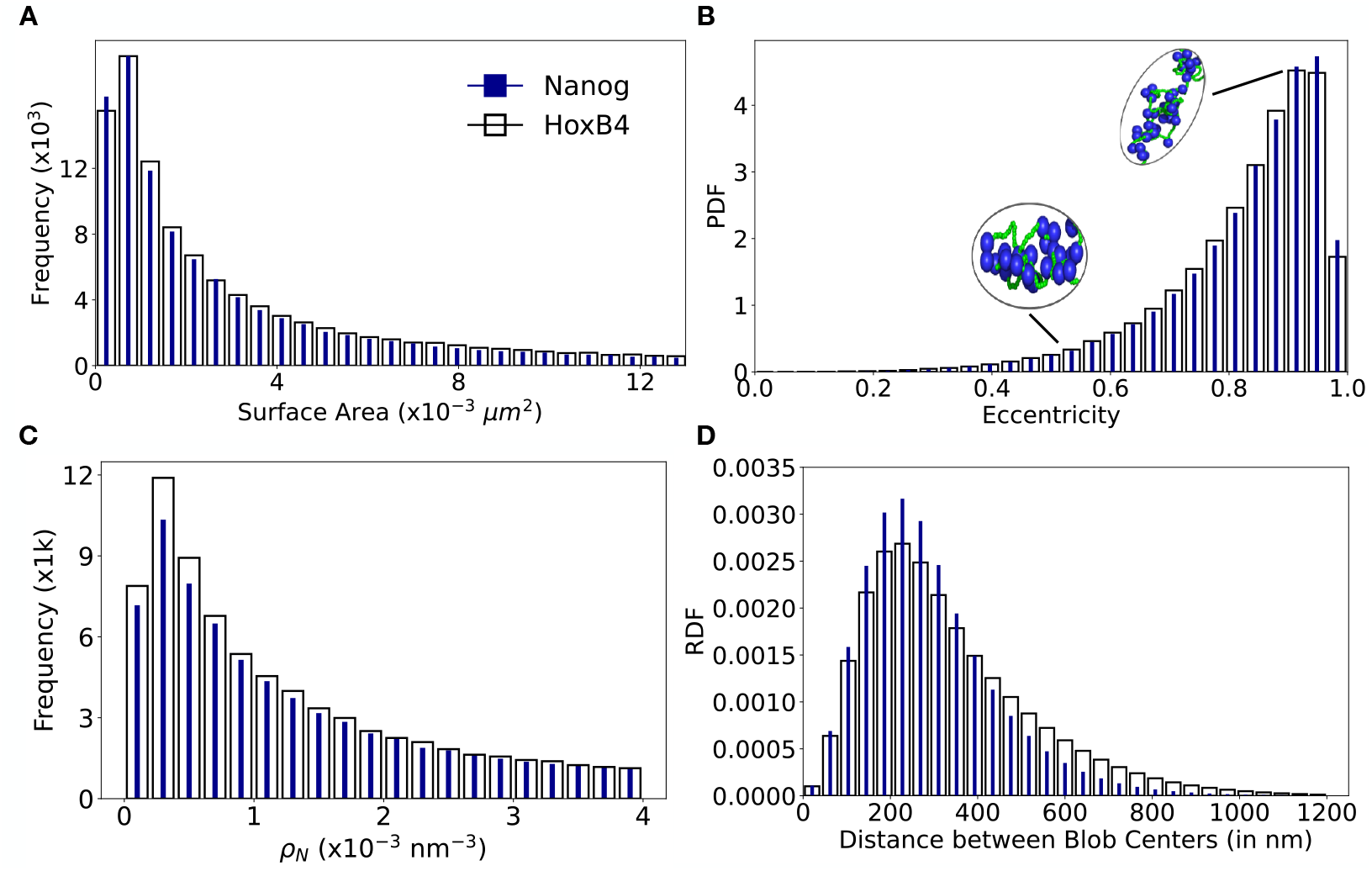
Characterizing the morphology of nucleosome blobs. A total of 13,000 NL-level chromatin conformations were analyzed to identify nucleosome blobs for both the Nanog (filled blue bars) and HoxB4 (open black bars) genomic loci. (A) shows the surface area distribution of blobs, estimated using the convex hull method across all blobs detected by DBSCAN. The results exhibit a lognormal fit with parameters 6.9 × 10^−3^ ± 0.019 *µm*^2^ for Nanog blobs and 7.1 × 10^−3^ ± 0.018 *µm*^2^ for HoxB4 blobs. (B) presents the eccentricity profile of the blobs, revealing their predominantly ellipsoidal shape. Representative snapshots of nucleosome blobs are shown in the inset. (C) depicts the nucleosome packing density profile for both Nanog and HoxB4 loci. (D) illustrates the spatial distribution of nucleosome blobs by estimating the radial distribution function (RDF) for both genomic loci.

How do these blobs look? To understand their shape, we performed Principal Component Analysis (PCA) on the convex hull of the set of beads involved in blob formation, which defines the boundary points of the blob’s shape. The first principal component corresponds to the major axis, and the second and third components correspond to the minor axes (or directions orthogonal to the major axis). To quantify the eccentricity of the convex hull, we computed the eccentricity of the best-fitting ellipsoid, derived from the relative lengths of the principal axes. A near-equal distribution of axis lengths is indicative of a spherical shape, whereas a pronounced disparity, particularly an elongation along one axis, suggests an ellipsoidal morphology. Thus, the eccentricity parameter provides a quantitative measure for assessing the geometric anisotropy of blobs comprising multiple nucleosomes. Our results, presented in Fig. 5B, reveals that the majority of blobs exhibit an elongated, ellipsoidal morphology, with their eccentricity predominantly concentrated near 1, indicating significant deviation from a spherical shape. The eccentricity profile peaks at 0.95 for Nanog blobs and at 0.91 for HoxB4 blobs, with average elongation of 43.7 nm and a width of 23.9 nm for the HoxB4 genomic loci, whereas slightly smaller blobs are observed in Nanog, with an elongation of approximately 41.8 nm and a width of 22.9 nm. To this end, we note that an experimental study based on super-resolution imaging of live human bone osteosarcoma (U2OS) cells reported a very similar eccentricity profile for chromatin blobs with most probable eccentricity of 0.9[64].

Having seen that most of the blobs in Nanog are slightly smaller than that of and HoxB4, we next examine their packing density (*ρ*_*N*_), quantified by the nucleosome density of each blob. Fig.5C presents the distribution of *ρ*_*N*_ for the chromatin blobs, revealing that the average packing density of HoxB4 blobs is approximately two times higher (0.032 ± 1.5 *nm*^−3^) than that of Nanog (0.015 ± 0.08 *nm*^−3^) blobs, delineating differences in their packing density. High fluctuations in the packing density however, is expected given the large variability in blob size distribution. We also probed if there is any organisational differences in the blobs between these two genomic loci. The results are presented in Fig. 5D, that shows a relatively a wider distribution of blobs in HoxB4 with parameters 333.5 ± 193.3 (mean ± sd) compared to that in Nanog (290.8 ± 152.9 (mean ± sd)) genomic loci. Furthermore, the shape of the distribution indicates a non-random and asymmetric distribution of blobs, favoring compact clustering with occasional larger separations, which is a characteristic feature of hierarchical organisation.

### 2.5 Differential packing density between Nanog and HoxB4 blobs results from differences in the nucleosome - nucleosome crosstalk

Having seen that the nucleosomes in most of the Nanog blobs are relatively loosely clustered (high packing density) compared to that in HoxB4 blobs, we pose the question that what makes HoxB4 blobs more compact than Nanog blobs? To address the issue, we constructed tetra-nucleosome contact maps for all consecutive sets of four nucleosomes along the chromatin segment of both genomic loci. Two nucleosomes were considered to be in contact if their inter-bead distance was less than 2.5 times of their diameter, consistent with the threshold used to generate the Micro-C contact map (see Materials and Methods). Given that the Nanog locus contains 998 nucleosomes and the HoxB4 segment consists of 1,036 nucleosomes, we constructed ensemble of tetra-nucleosome contact maps for both genomic loci, capturing their organizational variations across 13,000 conformations. The contact maps were then analyzed using k-means clustering to identify and classify distinct interaction patterns among nucleosomes within each genomic locus. A total of 10 clusters of tetra-nucleosome contact maps were generated, among which the three most significant clusters are presented in Fig. 6A-F. Fig. 6A-C denotes the significant tetra-nucleosome contact maps obtained for Nanog blobs. The first one (Fig. 6A) suggests that all four nucleosomes are widely spaced, characterizing an open arrangement of the nucleosomes (see the respective snap). The second and third tetra-nucleosome contact maps presented in Fig. 6B and 6C illustrate pairwise nucleosome interactions. Fig 6B presents interactions of the two central nucleosomes present in the middle of a tetra-nucleosome segment, suggesting a partially open conformation with localized compaction of nucleosomes in the middle. In contrast, Fig. 6C captures pairwise interactions between nucleosomes at both ends, forming two independent nucleosome pairs but no crosstalk between them. In contrast to Nanog, the tetra-nucleosome contact maps of HoxB4 blobs exhibit a markedly different organizational pattern, as shown in Fig. 6D–F. While the contact map in Fig. 6D still suggests a predominantly open arrangement of nucleosomes, it differs from Nanog in that adjacent nucleosomes occasionally engage in crosstalk, indicating a modest increase in the local compaction. The most striking difference, however, is observed in Fig. 6F, where all four nucleosomes engage in tight interactions, forming a fully closed state, that was not observed in Nanog blobs. A 31.56% fraction of the total cluster population is occupied by these closed-state tetra-nucleosome motifs, contributing to the formation of denser and more compact HoxB4 blobs compared to their Nanog counterparts.

**Fig. 6.**
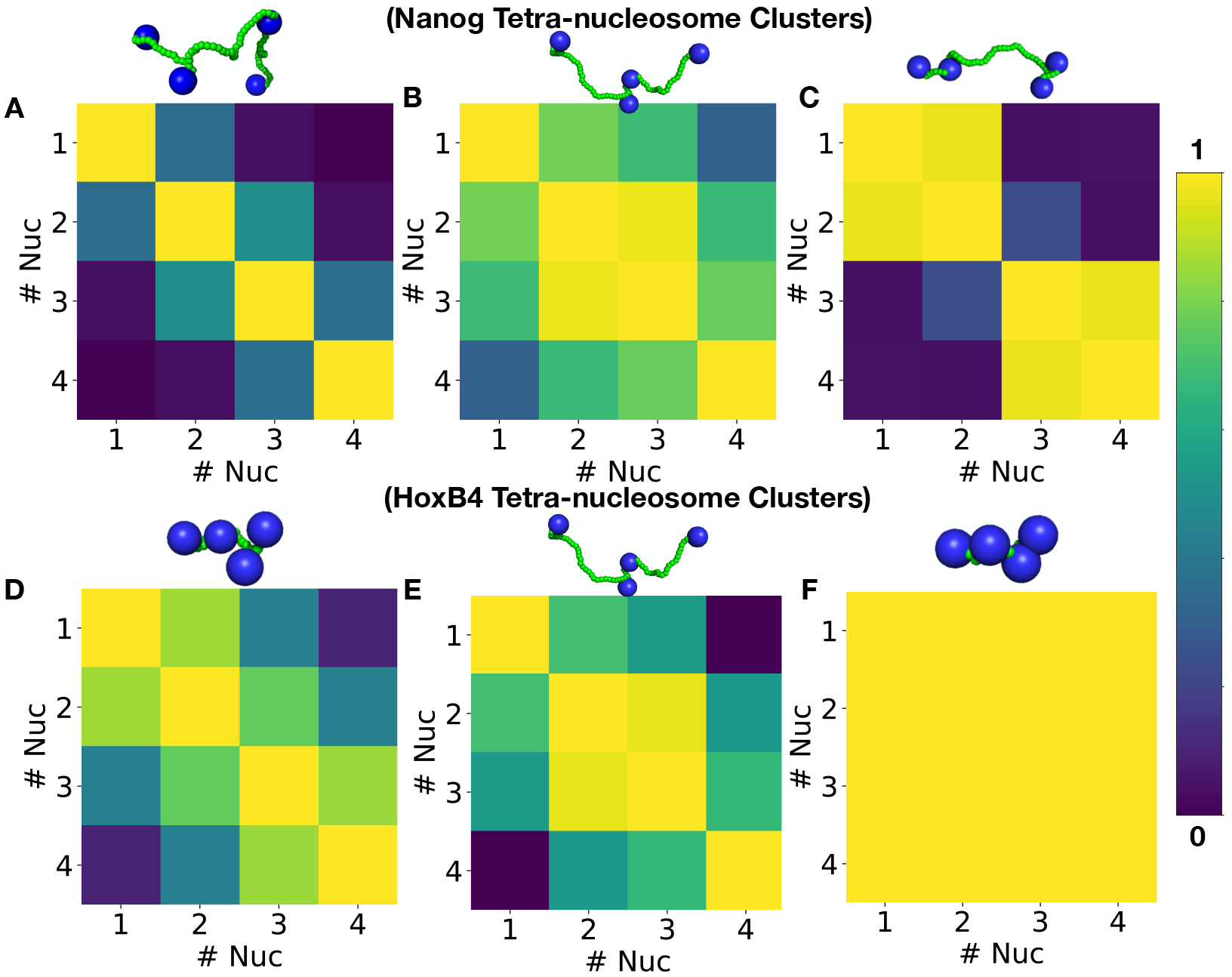
Tetra-nucleosome contact maps of Nanog and HoxB4 genomic loci. An ensemble of tetra-nucleosome contact maps was generated by considering all consecutive sets of four nucleosomes and identifying whether they form contacts. A contact was defined when the pairwise distance between two nucleosomes was within a cutoff distance *r*_*c*_, set as 2.5 times the nucleosome diameter. The k-means clustering algorithm was employed to identify the top three dominant clusters for both Nanog and HoxB4 genomic loci. Notably, these three clusters were the only ones comprising more than 10% of the total population of contact maps. (A–C) illustrate the significant tetra-nucleosome contact patterns observed for Nanog, while (D–F) depict the corresponding patterns for HoxB4. Representative structures of each contact map are shown above their respective maps. (A, D) indicate lower contact priority between adjacent nucleosomes, whereas (B, E) reveal dominant tetra-nucleosome interaction patterns, where the middle nucleosomes interact more frequently. In (C), interactions between i, i+1 and i+2, i+3 nucleosomes are predominant, while (F) displays a compact nucleosome arrangement, where multiple nucleosomes engage in interactions, suggesting a highly condensed chromatin structure.

### 2.6 Variation in blob distribution determines the accessibility of the genomic loci

Our second major finding highlights the differences in the radial distribution function (RDF) of nucleosome blob organization between Nanog and HoxB4, as shown in Fig. 5D. To investigate the origin of these differences, we further estimated the average blob size of participating nucleosomes. This averaging was performed over the same chromatin conformations generated from our simulations at NL-bead level resolution. The resulting distributions, presented in Fig. 7A–B, reveal two key aspects of nucleosome blob organization in Nanog and HoxB4 genomic loci: (1) The average blob size profile of nucleosomes exhibits a fluctuating behaviour (high standard deviation), suggesting blob formation an extremely dynamic process. Notably, 250 nm emerges as the critical distance at which blob motion significantly influences surface area. Beyond this threshold, the effect diminishes, suggesting that highly dynamic blobs are more spatially dispersed, whereas compact ones exhibit limited movement (see Fig. S7). Furthermore, Fig. S8 indicates that most blobs arise from the clustering of sequential nucleosomes, while only substantially large blobs encompass spatially proximate yet sequentially distant nucleosomes. (2) Fig. 7A-B also delineates the positions of the Nanog and HoxB4 genes within the selected chromatin regions, revealing a striking contrast in the local organization of nucleosome blobs. The average blob sizes in these regions indicate that Nanog is associated with significantly smaller blobs, comprising 40–80 nucleosomes, while HoxB4 exhibits larger blobs, containing 60–100 nucleosomes. However, it is noteworthy that at chromatin regions distal to the Nanog gene, much larger blobs emerge, surpassing those observed in HoxB4. This raises an intriguing question: what are the implications of this preferential distribution of smaller blobs near the Nanog locus compared to HoxB4?

**Fig. 7.**
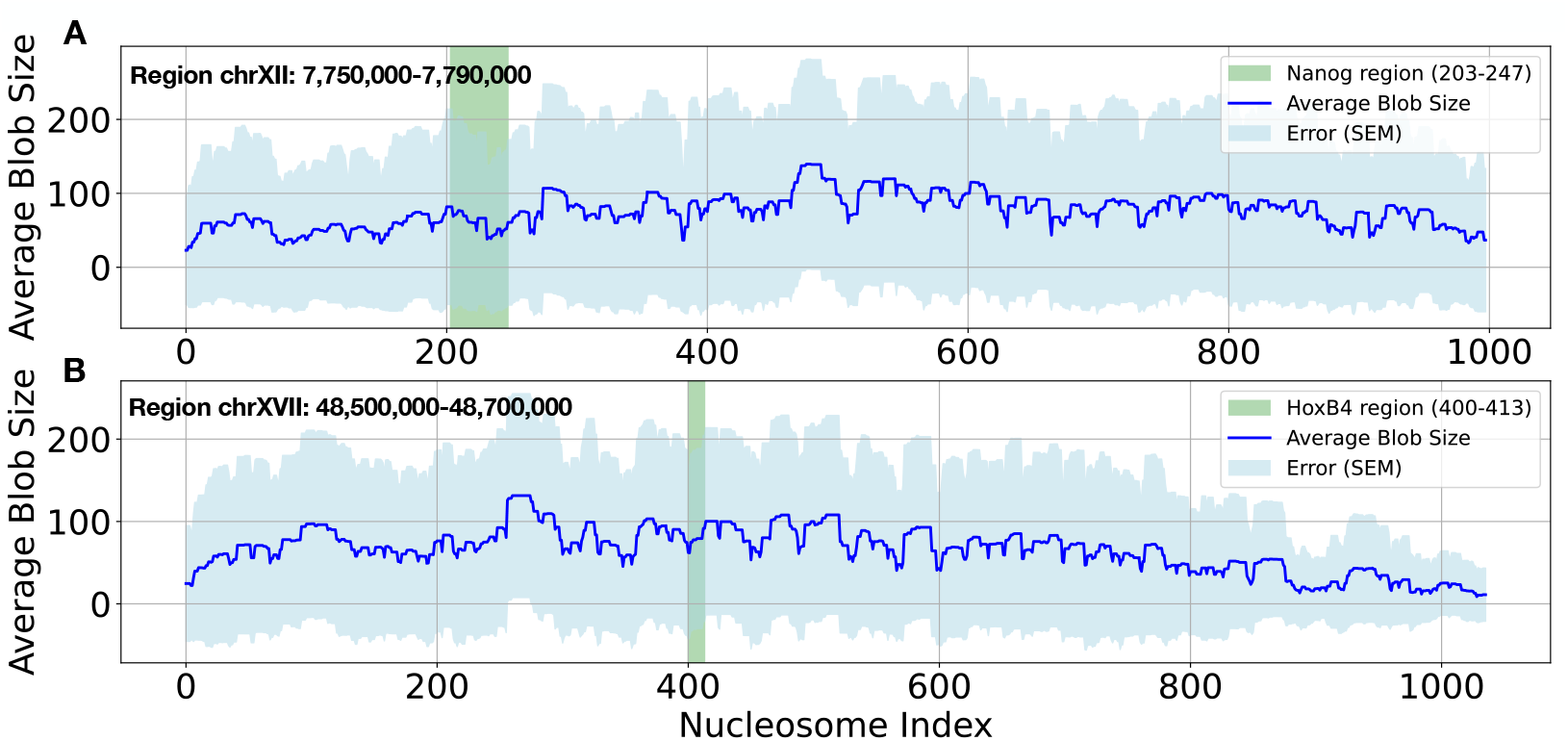
Average size of blobs formed by participating nucleosomes for Nanog (A) and HoxB4 (B) genomic loci. The green region highlights the location of the respective genes. A large deviation in blob sizes formed by any given nucleosome suggests a highly dynamic blob formation process, indicating variability in nucleosome clustering and chromatin organization.

To address this issue, we analyzed the nuclear environment of two genomic loci by computing covariance matrices from the three-dimensional Cartesian coordinates of nucleosome beads across 13,000 chromatin conformations per locus. We then performed Principal Component Analysis (PCA) on these covariance matrices and selected the two most significant components. To characterize the conformational landscape, we constructed free energy surfaces (FES) *F* (*PC*_1_, *PC*_2_) defined as *F* (*PC*_1_, *PC*_2_) = −*k*_*B*_*T* ln *P* (*PC*_1_, *PC*_2_), where *P* (*PC*_1_, *PC*_2_) represents the probability of the system occupying a given state in terms of the selected components. These free energy surfaces provide a quantitative representation of chromatin accessibility, structural flexibility, and thermodynamic stability of the genomic locus within its nuclear environment.

The results in Fig. 8A-B reveal significant differences in the local chromatin environment surrounding the Nanog and HoxB4 genomic loci. The free energy landscape of the Nanog locus (Fig. 8A) is depicted as a function of its two most dominant PCA components, which together account for approximately 71% of the total variance—substantially more than the remaining components. The resulting free energy surface is highly rugged, characterized by multiple low-energy basins, indicating that the locus explores a broader conformational space. In contrast, Fig. 8B illustrates the free energy landscape of the HoxB4 locus, also represented using its two most significant PCA components, which capture roughly 74% of the total variance. Unlike Nanog, the HoxB4 free energy surface exhibits fewer minima and appears more constrained, suggesting a more restricted conformational ensemble. Further analysis of the free energy surfaces reveals 2,609 low-energy basins for Nanog, significantly more than the 1,688 observed for HoxB4. This suggests that Nanog explores a larger number of stable conformations and is more dynamic. To quantify this conformational diversity, we computed the Boltzmann-weighted entropy, given by, *S* = −*k*_*B*_ ∑ *P* (*PC*_1_, *PC*_2_) ln *P* (*PC*_1_, *PC*_2_) where ∑ *P* (*PC*_1_, *PC*_2_) = 1 represents the normalized Boltzmann probability of a given state. Our results show that HoxB4 has a Boltzmann-weighted entropy of 6.99 *k*_*B*_, significantly lower than that of Nanog (7.39 *k*_*B*_), further confirming that Nanog samples a broader range of conformational states, supporting its greater dynamical flexibility. The lower entropy of HoxB4 aligns with its constrained free energy landscape, indicating that it likely remains in a few specific conformations. Thus, our combined PCA-based free energy landscapes and Boltzmann-weighted entropy analyses suggest that Nanog exhibits greater conformational flexibility, making it more accessible and dynamically adaptable. In contrast, HoxB4 appears structurally constrained, implying a more stable, functionally rigid state.

**Fig. 8.**
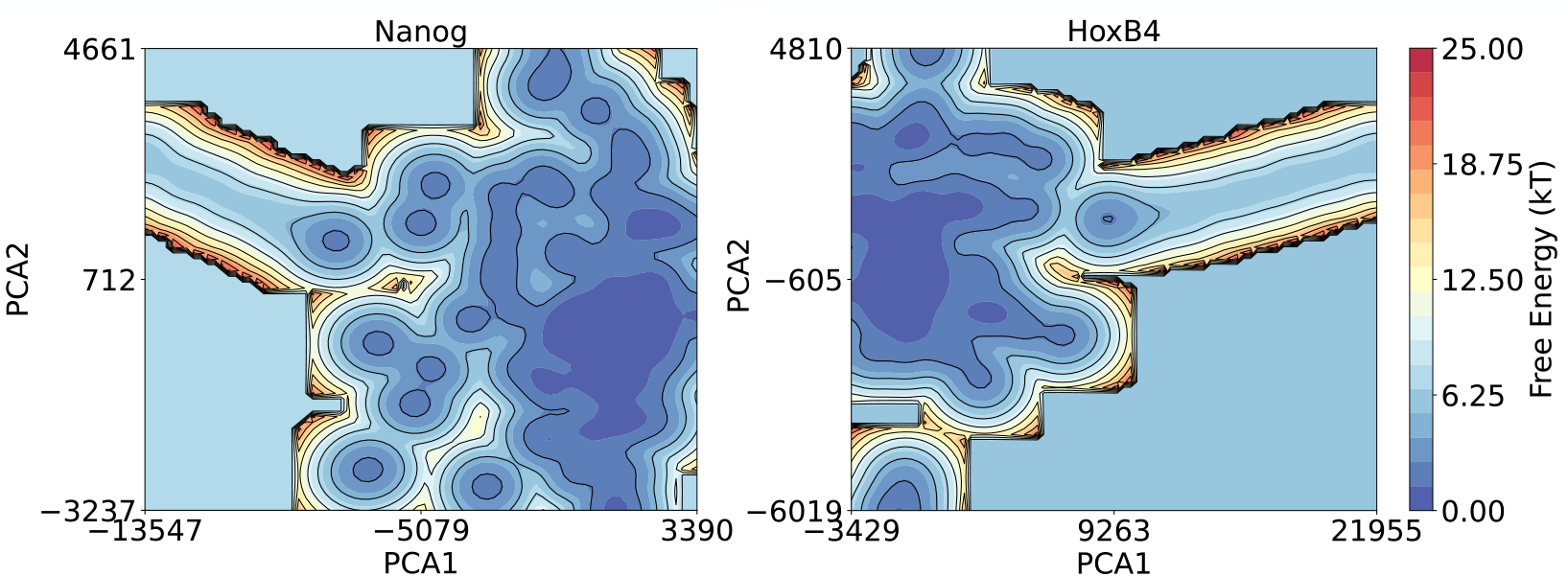
Free Energy Surfaces of Nanog and HoxB4. (A) Free energy surface of Nanog, represented by its two most dominant PCA components (71% variance). The rugged landscape with multiple low-energy basins (blue) indicates high conformational flexibility. (B) Free energy surface of HoxB4, based on its top two PCA components (74% variance). Compared to Nanog, HoxB4 has fewer minima and a more constrained landscape, suggesting restricted conformational dynamics. Color bars indicate free energy in *k*_*B*_*T*, with blue representing stable conformations and red indicating higherenergy states.

To further substantiate our claim, we quantified three additional parameters to characterize the local nuclear environment surrounding the two genomic loci: (i) the mean inter-nucleosomal distance, *< R*_*nuc*_ *>*, among interacting nucleosomes, (ii) the number of interacting nucleosomes, *N*_*nuc*_ and (iii) radius of gyration (*ROG*). To focus on the local chromatin environment, we analyzed chromatin segments spanning 50 nucleosomes upstream and downstream of each locus. The results presented in Fig. S9, S10 and S11 (Supplementary Text) reveal that the mean *< R*_*nuc*_ *>* for the Nanog locus is 123.7 ± 11.8, whereas for the HoxB4 locus, it is 113.5 ± 11.6, indicating that nucleosomes are more densely packed around HoxB4 compared to Nanog. The most probable number of neighboring nucleosomes (*N*_*nuc*_) within a 25 nm contact distance is approximately 3 for the Nanog locus, whereas for HoxB4, it is around 4. Additionally, the distribution of the radius of gyration for the local chromatin segments encompassing each genomic locus shows that the *< ROG >* for the Nanog locus is 96.9 ± 10.4, while for HoxB4, it is 89.1 ± 10.1. This suggests that the local chromatin environment surrounding HoxB4 is significantly more constrained and compact compared to Nanog, reinforcing the notion of a structurally restricted nuclear architecture for HoxB4.

The stark contrast between the two free energy surfaces, along with the supporting analyses (also see Supplementary Text) therefore, strongly suggests that Nanog locus, with its rugged, shallow multiple low-energy basins, resides in a more flexible and open chromatin region, making it highly accessible for molecular interactions. In contrast, HoxB4 appears to be buried within a dense chromatin cluster, rendering it less accessible due to its pronounced energy well and steeper energy barriers. These findings indicate that Nanog’s accessibility may facilitate regulatory interactions and transcriptional activity, whereas HoxB4’s confinement likely contributes to its repressed state, potentially functioning as part of a structurally stabilized or transcriptionally silenced chromatin domain.

### 2.7 Segment wise variation in chromatin rigidity driven by nucleosome blob organization

We also investigated how variations in nucleosome blob size influence the overall chromatin architecture. The chromatin segments involved in the formation of the blob contribute to the persistence of the chromatin chain along the elongation axis of the blob. Consequently, differences in blob size and compactness are expected to modulate the rigidity of distinct chromatin regions. To examine the variation in chromatin bending rigidity across different regions, we employed a rolling window approach with a window size of 1,500 NL-beads and an overlap of 500 beads. Within each window, we estimated the persistence length by computing the cosine of the angles between bond vectors connecting adjacent NL-beads. The bending rigidity was then derived from the segment-wise persistence length, and the results for the Nanog and HoxB4 genomic loci are presented in Fig. 9A. Our results clearly demonstrate that the Nanog locus exhibits significantly lower bending rigidity compared to the HoxB4 locus, indicating a more flexible chromatin chain. This increased flexibility may facilitate the formation of long-range contacts between distant chromatin segments, potentially influencing higher-order chromatin organization and transcriptional regulation.

**Fig. 9.**
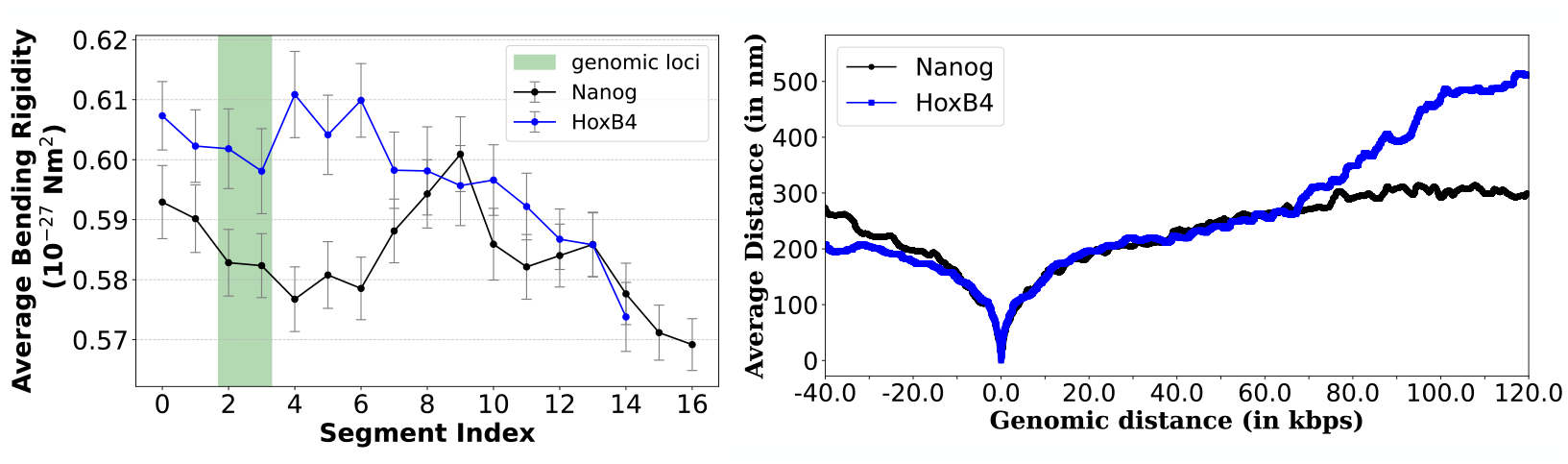
Segment-wise variational bending rigidity of chromatin fibers encompassing Nanog and HoxB4 gene loci and its impact on long-range genomic interactions. (A) presents the bending rigidity of chromatin stretches associated with Nanog and HoxB4 loci, estimated from the segment-wise persistence length of the chromatin fiber. Variations in bending rigidity reflect differences in the propensity of segments to form nucleosome blobs. (B) illustrates how these differences in bending rigidity influence long-range genomic interactions by estimating the distances of genomic segments from the transcription initiation sites (TIS) of the two genes. The relatively larger distance of downstream genomic segments from the TIS in HoxB4 suggests a reduced likelihood of long-range enhancer-promoter interactions, in contrast to the Nanog locus, where such interactions are more probable.

To test our hypothesis, we measured the spatial distances of all chromatin segments from the transcription start site (TSS) of the Nanog and HoxB4 loci. As shown in Fig. 9, while the TSS and promoter regions of both genes remain in close proximity (∼200*nm*), the distance between the TSS and downstream chromatin segments increases progressively in HoxB4 as the sequential gap grows. In contrast, at the Nanog locus, the TSS remains at a relatively constant spatial distance from downstream chromatin segments over a long stretch (∼160*kbps*). This suggests that long-range interactions between downstream enhancers and the TSS-promoter region are structurally feasible in Nanog, whereas the higher rigidity of the HoxB4 chromatin chain prevents such interactions due to increased spatial separation. Notably, unlike Nanog in hESCs, no downstream enhancer has been reported for HoxB4, further aligning with our findings.

## 3 Methods

Our experimentally informed polymer model for capturing chromatin organization at multiple resolutions follows a two-tiered approach. At the first level, we utilize Hi-C contact maps at a 5 kb resolution to reconstruct the global architecture of a 0.2 Mb chromatin segment, though the method is not inherently restricted to a specific length. However, as the chromatin segment length increases, computational complexity scales exponentially. A Hi-C contact map represents an ensemble-averaged contact frequency matrix derived from a population of cells. Each contact in a Hi-C map reflects the probability of two genomic segments being spatially close across many different chromatin conformations rather than a single definitive structure. Consequently, chromatin function is not dictated by a single conformation but rather by an ensemble of possible structures. A plausible approach to capturing those representative chromatin conformations from the Hi-C map is to decompose it into an ensemble of contact maps, ensuring that their averaged contact frequencies reproduce the original Hi-C contact map. To this end, it is noteworthy that not all contacts in a Hi-C map are equally significant. While some contacts occur frequently and play a crucial role in chromatin organization, a substantial number of contacts form only sporadically, as reflected in their lower contact frequencies. Distinguishing between these persistent and transient interactions is essential for accurately reconstructing chromatin architecture and understanding its functional implications.

To identify the persistent contacts crucial for chromatin organization, we analyzed the KR-normalized Hi-C contact map of the target chromatin segment at a 5 kb resolution. We selected only those contacts whose probabilities exceeded a threshold specific to the probability distribution at each genomic separation distance (|*j* −*i*|). The threshold was defined as the sum of the mean and standard deviation of the probability distribution for each (|*j* – *i*|) value, ensuring that only significantly frequent contacts were retained for further analysis [65]. The resulting matrix of important contacts is then stochastically decomposed into an ensemble of contact matrices (see Supplementary Text for details). Simulations are subsequently performed on a homopolymer model, with each bead representing 5kbps, folding the polymer according to these individual contact matrices to generate an ensemble of chromatin conformations. The energetics of these structures are governed by (see Supplementary text for more details):

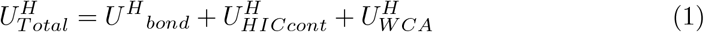

where, (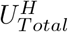 represents the total potential energy of the homopolymer chain. The Weeks-Chandler-Andersen (WCA) potential, 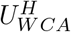, accounts for steric interactions and prevents the overlap of genomic segmental beads. The bonding potential, 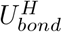, ensures connectivity between adjacent beads. Additionally, non-adjacent genomic pairs that are in contact according to the Hi-C contact map are coupled via 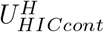, in which case 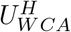 does not apply to these specific pairs.

In the second stage, the chromatin fiber is modeled as a copolymer, with two distinct bead types representing nucleosomes and linker DNA segments. The nucleosome beads, which are larger, correspond to 142 base pairs, while the smaller linker DNA beads represent approximately 8 base pairs (see Supplementary Text for details). This approach is similar to the one proposed earlier by Wiese [69]. The placement of nucleosome beads along the copolymer chain is determined using MNase-Seq experimental data, analyzed with the help of bioinformatics pipeline DANPOS [79]. In this experiment, Micrococcal Nuclease (MNase) selectively digests unprotected DNA, leaving behind nucleosome-bound fragments that are then aligned to the genome, providing a high-resolution map of nucleosome positions. The copolymer model is subsequently simulated under the following energetic constraints until it satisfies the individual contact maps derived from the homopolymer structures obtained in the previous step.

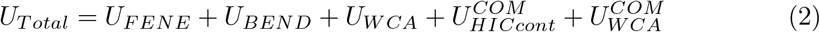

Where, *U*_*T otal*_ represents the total energy of the copolymer chain (see Supplementary text for more details). The finite extensible nonlinear elastic (FENE) potential, *U*_*F ENE*_, acts as a bonding force between adjacent beads, maintaining chain connectivity. The bending potential, *U*_*BEND*_, governs the angular constraints imposed by three consecutive linker DNA beads within the copolymer chain, ensuring realistic flexibility. The Weeks-Chandler-Andersen (WCA) potential, *U*_*W CA*_, accounts for steric interactions and prevents overlap between non-adjacent beads in the copolymer chain. We coarse-grained the NL beads corresponding to every bead in the homopolymer chain and introduced a bonding potential between the center-of-mass positions of the NL bead segments, represented by 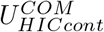, to ensure the contacts observed in the homopolymer chain. The steric repulsion is enforced through 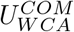 to prevent overlap.

Once all contacts are satisfied, the simulations are extended up to 6,500,000 steps to generate an ensemble of high-resolution chromatin conformations at the nucleosome-linker (NL) bead resolution, with structures recorded at every 50,000 MD steps. The resultant conformations thus, not only provide a near base-pair resolution description but also faithfully preserve the large-scale chromatin organization captured by the Hi-C contact map at a 5 kb resolution.

To this end, it is important to note that even for a 0.2 Mb chromatin segment, the NL-bead representation results in a substantially large system size for simulations. To mitigate the associated computational challenges, we performed our simulations on a GPU using a custom-developed code written in CUDA C, utilizing CUDA version 11.0 and the GeForce GTX 10 architecture. This approach provides a 25 speedup compared to running the equivalent code on a CPU for the current system size. The simulations employ Langevin dynamics, where the velocity-Verlet algorithm is used for time integration with a time step of Δ*t* = 0.005*τ*, where *τ* represents the fundamental simulation time unit set at 4. All simulations are conducted at *k*_*B*_*T* = 1, with all energy functions and *τ* expressed in dimensionless units.

## 4 Conclusion

By amalgamating two distinct polymer models that describe chromatin conformations at different scales, we present a method to bridge multiscale chromatin organization, enabling the capture of both local and global chromatin structures at a reasonably high resolution. Our method demonstrates chromatin fibre as an array of nucleosomes and linker DNA (NL-representation), where each linker bead represents around 8 bps. The advantage of this representation is that it enables the visualization of genomic loci at near base-pair resolution, facilitating the investigation of their involvement in crosstalk with spatially proximal genes. Building on this concept, we applied our method to three distinct 0.2 Mbps chromatin segments from human chromosomes VII, XII, and XVII. We demonstrate that our model effectively reconstructs chromatin conformations, with the ensemble-averaged contact maps closely matching the experimental Micro-C and Hi-C contact maps. The strong agreement between simulated and experimental contact frequencies, as evidenced by high Pearson and Spearman correlation coefficients, underscores the robustness of our approach in modeling chromatin architecture.

A key insight from our analysis is the emergence of spatially heterogeneous nucleosome “blobs” as fundamental organizational units of chromatin. These blobs exhibit nonrandom clustering patterns, with significant correlations between intra-blob nucleosome interactions and Hi-C contact maps. Notably, our model also identifies sub-TAD domains that remain undetectable at lower Hi-C resolutions, providing a finer-scale perspective on chromatin folding dynamics. Further analysis of these nucleosome blobs reveals striking similarities to the morphology of blob-like structures observed in superresolution imaging of chromatin in living human cells. Most significantly, our findings demonstrate a lognormal distribution of blob sizes that closely mirrors experimental observations [64] but sharply contradicts several computational studies that describe chromatin as a crumpled globule polymer [55, 56].

Leveraging our approach, we also distinguish between the nuclear environments of transcriptionally active (Nanog) and repressive (HoxB4) genomic loci. Our results show that HoxB4 blobs exhibit a more compact organization with higher nucleosome-nucleosome interactions, whereas Nanog blobs are more loosely clustered, displaying greater variability in size and shape. This difference in packing density is directly linked to variations in tetra-nucleosome interactions, with HoxB4 showing a greater propensity for closed nucleosome arrangements. On a larger scale (0.2 Mbps), our analysis reveals a striking contrast between the chromatin environments of Nanog and HoxB4. The Nanog locus exhibits a highly rugged free energy landscape with numerous low-energy basins, higher entropy, and greater conformational flexibility, indicating an open and dynamic chromatin state. In contrast, HoxB4 displays a more constrained, less rugged free energy landscape, lower entropy, and a compact chromatin environment, suggesting a structurally restricted and less accessible state. These findings imply that Nanog is more amenable to regulatory interactions and transcriptional activity, while HoxB4 likely resides in a repressed or structurally stabilized chromatin domain. Moreover, we reveal that the distribution of nucleosome blobs imparts varying degrees of rigidity across different regions of the chromatin fiber, leading to overall higher stiffness around the HoxB4 locus compared to Nanog. This increased stiffness significantly reduces the probability of long-range enhancer-promoter interactions in HoxB4 compared to the Nanog genomic locus, providing a mechanistic explanation for the differences in their transcriptional activities observed experimentally in hESCs.

To this end, it is important to note that our model operates without any adjustable parameters and relies entirely on experimental inputs. Consequently, the quality of the input data plays a crucial role in the accuracy of our predictions. While we have demonstrated that our model performs well using ensemble-averaged Hi-C and nucleosome positioning data, we anticipate that applying it to single-cell data would provide a more detailed structural representation, thereby enabling a more precise link to functional roles. Furthermore, our current model assumes nonspecific short-range interactions among spatially proximal nucleosomes. However, a more specific and fine-tuned internucleosomal potential is necessary to accurately predict the impact of epigenetic modifications on chromatin folding. Developing such a refined potential remains a key objective for our future work. Despite these limitations, our findings establish nucleosome blobs as fundamental components of chromatin architecture and highlight how their spatial organization influences nuclear environment of genomic loci and thereby their functions. By elucidating chromatin structures across multiple resolutions, our study lays the groundwork for future research into the intricate relationships between chromatin organization, epigenetic modifications, and transcriptional regulation in normal and diseased cell lines.

## Supplementary information

The supplementary text integrates detailed model information with a comprehensive description of the methods. Additionally, it includes data exportation details for the Hi-C micro-contact map and nucleosome positions.

## Acknowledgements

We gratefully acknowledge the financial support from DST India (CRG/2023/000636), DBT India (BT/PR46247/BID/7/1015/2023) and DBT CoE research grant. A.B gratefully acknowledges support from the Alexandar von Humboldt Foundation, Germany. D.H gratefully acknowledges the support from Deutsche Forschungsgemeinschaft (DFG, German Research Foundation) under Germany’s Excellence Strategy EXC 2181/1 - 390900948 (the Heidelberg STRUCTURES Excellence Cluster). R.M acknowledges the financial support from the Council of Scientific & Industrial Research (CSIR), Govt. of India for Senior Research Fellowship (File No: 09/0263(11815)/2021-EMR-I).

## Declarations

The authors declare no competing interests.

## References

[1] Forth, S., et al.: Torque measurement at the single-molecule level. Annual Review of Biophysics 42, 583–604 (2013)

[2] Ma, J., et al.: Dna supercoiling during transcription. Biophysical Reviews 8(Suppl 1), 75–87 (2016)

[3] Dunaway, M., et al.: Local domains of supercoiling activate a eukaryotic promoter in vivo. Nature 361(6414), 746–748 (1993)

[4] Mondal, A., et al.: Understanding the role of dna topology in target search dynamics of proteins. J. Phys. Chem. B 121(40), 9372–9381 (2017)

[5] Bhattacherjee, A., et al.: Search by proteins for their dna target site: 1. the effect of dna conformation on protein sliding. Nucleic Acids Res 42(20), 12404–12414 (2014)

[6] Bhattacherjee, A., et al.: Search by proteins for their dna target site: 2. the effect of dna conformation on the dynamics of multidomain proteins. Nucleic Acids Res 42(20), 12415–12424 (2014)

[7] Wang, X., et al.: Negatively charged, intrinsically disordered regions can accelerate target search by dna-binding proteins. Nucleic Acids Res 51(10), 4701–4712 (2023)

[8] Vuzman, D., et al.: Dna search efficiency is modulated by charge composition and distribution in the intrinsically disordered tail. Proceedings of the National Academy of Sciences 107(49), 21004–21009 (2010)

[9] Sangeeta, et al.: Role of shape deformation of dna-binding sites in regulating the efficiency and specificity in their recognition by dna-binding proteins. JACS Au 4(7), 2640–2655 (2024)

[10] Sangeeta, et al.: Nick induced dynamics in supercoiled dna facilitates the protein target search process. J. Phys. Chem. B 128(34), 8246–8258 (2024)

[11] Mondal, A., et al.: Torsional behaviour of supercoiled dna regulates recognition of architectural protein fis on minicircle dna. Nucleic Acids Res 50(12), 6671–6686 (2022)

[12] Dey, P., others.: Structural basis of enhanced facilitated diffusion of dna-binding protein in crowded cellular milieu. Biophysical Journal 118(2), 505–517 (2020)

[13] Dey, P., et al.: Role of macromolecular crowding on the intracellular diffusion of dna binding proteins. Scientific Reports 8(1), 844 (2018)

[14] Mishra, S., et al.: How do nucleosome dynamics regulate protein search on dna? The Journal of Physical Chemistry B 127(25), 5702–5717 (2023)

[15] Tsompana, M., et al.: Chromatin accessibility: a window into the genome. Epigenetics Chromatin 7(1), 33 (2014)

[16] Hofmann, A., et al.: Self-organised segregation of bacterial chromosomal origins. eLife 8, 46564 (2019)

[17] Mondal, A., et al.: Nucleosome breathing facilitates cooperative binding of pluripotency factors sox2 and oct4 to dna. Biophysical Journal 121(23), 4526– 4542 (2022)

[18] Bilokapic, S., et al.: Histone octamer rearranges to adapt to dna unwrapping. Nature Structural & Molecular Biology 25(1), 101–108 (2018)

[19] Lee, B., et al.: Characterizing chromatin interactions of regulatory elements and nucleosome positions, using hi-c, micro-c, and promoter capture micro-c. Epigenetics & Chromatin 15(1), 41 (2022)

[20] Fraser, J., et al.: Hierarchical folding and reorganization of chromosomes are linked to transcriptional changes in cellular differentiation. Molecular Systems Biology 11(12), 852 (2015)

[21] Luger, K., et al.: Crystal structure of the nucleosome core particle at 2.8 Å resolution. Nature 389(6648), 251–260 (1997)

[22] Mondal, A., et al.: Kinetic origin of nucleosome invasion by pioneer transcription factors. Biophysical Journal 120(23), 5219–5230 (2021)

[23] Fierz, B., et al.: Biophysics of chromatin dynamics. Annual Review of Biophysics 48, 321–345 (2019)

[24] Brandani, G., et al.: Dna sliding in nucleosomes via twist defect propagation revealed by molecular simulations. Nucleic Acids Research 46(6), 2788–2801 (2018)

[25] Meersseman, G., et al.: Mobile nucleosomes—a general behavior. The EMBO Journal 11(8), 2951–2959 (1992)

[26] Li, K., et al.: Inter-nucleosomal potentials from nucleosomal positioning data. The European Physical Journal E 45(4), 33 (2022)

[27] Mishra, S., et al.: Superstructure detection in nucleosome distribution shows common pattern within a chromosome and within the genome. Life 12(4), 541 (2022)

[28] Farr, S., et al.: Nucleosome plasticity is a critical element of chromatin liquid–liquid phase separation and multivalent nucleosome interactions. Nature Communications 12(1), 2883 (2021)

[29] Dekker, J., et al.: Capturing chromosome conformation. Science 295(5558), 1306– 1311 (2002)

[30] Bonev, B., et al.: Organization and function of the 3d genome. Nature Reviews Genetics 17(11), 661–678 (2016)

[31] Sati, S., et al.: Chromosome conformation capture technologies and their impact in understanding genome function. Chromosoma 126(1), 33–44 (2017)

[32] Lieberman-Aiden, E., et al.: Comprehensive mapping of long-range interactions reveals folding principles of the human genome. Science 326(5950), 289–293 (2009)

[33] Sanborn, A., et al.: Chromatin extrusion explains key features of loop and domain formation in wild-type and engineered genomes. Proceedings of the National Academy of Sciences 112(47), 6456–6465 (2015)

[34] Galazka, J., et al.: Neurospora chromosomes are organized by blocks of importin alpha-dependent heterochromatin that are largely independent of h3k9me3. Genome Research 26(8), 1069–1080 (2016)

[35] Shah, S., et al.: Dynamics and spatial genomics of the nascent transcriptome by intron seqfish. Cell 174(2), 363–376 (2018)

[36] Franke, M., et al.: Formation of new chromatin domains determines pathogenicity of genomic duplications. Nature 538(7624), 265–269 (2016)

[37] Flavahan, W., et al.: Insulator dysfunction and oncogene activation in idh mutant gliomas. Nature 529(7584), 110–114 (2016)

[38] Bonev, B., et al.: Multiscale 3d genome rewiring during mouse neural development. Cell 171(3), 557–57224 (2017)

[39] Lee, B., et al.: Characterizing chromatin interactions of regulatory elements and nucleosome positions, using Hi-C, Micro-C, and promoter capture Micro-C. Epigenetics & Chromatin 15, 41 (2022)

[40] Dixon, J., et al.: Topological domains in mammalian genomes identified by analysis of chromatin interactions. Nature 485(7398), 376–380 (2012)

[41] Zhang, Y., Heermann, D.W.: Loops determine the mechanical properties of mitotic chromosomes. PLOS ONE 6(12), 1–13 (2011)

[42] Pope, B., et al.: Topologically associating domains are stable units of replicationtiming regulation. Nature 515(7527), 402–405 (2014)

[43] Symmons, O., et al.: Functional and topological characteristics of mammalian regulatory domains. Genome Research 24 (2014)

[44] Ramírez, F., et al.: High-resolution tads reveal dna sequences underlying genome organization in flies. Nature Communications 9(1), 189 (2018)

[45] Lupiáñez, D., et al.: Disruptions of topological chromatin domains cause pathogenic rewiring of gene-enhancer interactions. Cell 161(5), 1012–1025 (2015)

[46] Lupiáñez, D., et al.: Breaking tads: How alterations of chromatin domains result in disease. Trends in Genetics 32(4), 225–237 (2016)

[47] Hovenga, V., Kalita, J., Oluwadare, O.: Hic-gnn: A generalizable model for 3d chromosome reconstruction using graph convolutional neural networks. Computational and Structural Biotechnology Journal 21, 812–836 (2023)

[48] Beagrie, R.A., Scialdone, A., et al.: Complex multi-enhancer contacts captured by genome architecture mapping. Nature 543(7646), 519–524 (2017)

[49] Kalhor, R., Tjong, H., et al.: Genome architectures revealed by tethered chromosome conformation capture and population-based modeling. Nature Biotechnology 30(1), 90–98 (2012)

[50] Hu, M., Deng, K., et al.: Bayesian inference of spatial organizations of chromosomes. PLOS Computational Biology 9(1), 1–14 (2013)

[51] Le Treut, G., Képès, F., Orland, H.: A polymer model for the quantitative reconstruction of chromosome architecture from hic and gam data. Biophysical Journal 115(12), 2286–2294 (2018)

[52] Di Pierro, M., et al.: De novo prediction of human chromosome structures: Epigenetic marking patterns encode genome architecture. Proceedings of the National Academy of Sciences of the United States of America 114(46), 12126–12131 (2017)

[53] Zhang, B., Wolynes, P.G.: Prediction of chromosome conformations with maximum entropy principle. Biophysical Journal 108(2), 537 (2015)

[54] Marti-Renom, M.A., Mirny, L.A.: Bridging the resolution gap in structural modeling of 3d genome organization. PLOS Computational Biology 7(7), 1–6 (2011)

[55] Mirny, L.A., et al.: The fractal globule as a model of chromatin architecture in the cell. Chromosome Res. 19(1), 37–51 (2011)

[56] Polovnikov, K.E., et al.: Crumpled polymer with loops recapitulates key features of chromosome organization. Phys. Rev. X 13(4), 041029 (2023)

[57] Shi, G., Thirumalai, D.: A maximum-entropy model to predict 3d structural ensembles of chromatin from pairwise distances with applications to interphase chromosomes and structural variants. Nature Communications 14(1), 1150 (2023)

[58] Lin, X., Qi, Y., Latham, A.P., Zhang, B.: Multiscale modeling of genome organization with maximum entropy optimization. The Journal of Chemical Physics 155(1), 010901 (2021)

[59] Nguyen, H.T., Thirumalai, D.: Liquid-liquid phase separation of repeat disorder sequences leads to rna conformational and dynamical heterogeneity. Biophysical Journal 120(3), 108 (2021)

[60] Zhang, B., Wolynes, P.G.: Topology, structures, and energy landscapes of human chromosomes. Proceedings of the National Academy of Sciences 112(19), 6062– 6067 (2015)

[61] Galan, S., Serra, F., et al.: Identification of chromatin loops from hi-c interaction matrices by ctcf–ctcf topology classification. NAR Genomics and Bioinformatics 4(1) (2022)

[62] Dugar, G., et al.: A chromosomal loop anchor mediates bacterial genome organization. Nature Genetics 54(2), 194–201 (2022)

[63] Hofmann, A., et al.: The role of loops on the order of eukaryotes and prokaryotes. FEBS Letters 589(20PartA), 2958–2965 (2015)

[64] Barth, R., et al.: Coupling chromatin structure and dynamics by live super-resolution imaging. Sci. Adv. 6(27), 2196 (2020)

[65] Kadam, S., et al.: Predicting scale-dependent chromatin polymer properties from systematic coarse-graining. Nature Communications 14(1), 4108 (2023)

[66] Lorzadeh, A., Bilenky, M., Hammond, C., Knapp, D., Li, L., Miller, P., Carles, A., Heravi-Moussavi, A., Gakkhar, S., Moksa, M., et al.: Nucleosome Density ChIP-Seq Identifies Distinct Chromatin Modification Signatures Associated with MNase Accessibility. Cell Reports 17, 2112–2124 (2016)

[67] Consortium, Roy, S., Ernst, J., Kharchenko, P., Kheradpour, P., Negre, N., Eaton, M., Landolin, J., Bristow, C., Ma, L., et al.: Identification of functional elements and regulatory circuits by Drosophila modENCODE. Science 330, 1787–1797 (2010)

[68] Rosenbloom, K., Sloan, C., Malladi, V., Dreszer, T., Learned, K., Kirkup, V., Wong, M., Maddren, M., Fang, R., Heitner, S., et al.: ENCODE data in the UCSC Genome Browser: year 5 update. Nucleic Acids Research 41, 56–63 (2013)

[69] Wiese, O., et al.: Nucleosome positions alone can be used to predict domains in yeast chromosomes. Proceedings of the National Academy of Sciences 116(35), 17307–17315 (2019)

[70] Ricci, M., et al.: Chromatin fibers are formed by heterogeneous groups of nucleosomes in vivo. Cell 160(6), 1145–1158 (2015)

[71] Consortium, R.E., et al.: Integrative analysis of 111 reference human epigenomes. Nature 518, 317–330 (2015)

[72] Ernst, J., Kellis, M.: Chromatin-state discovery and genome annotation with ChromHMM. Nature Protocols 12(12), 2478–2492 (2017)

[73] Roy, Ernst, Jason, et al.: Identification of functional elements and regulatory circuits by drosophila modencode. Science 330(6012), 1787–1797 (2010)

[74] Yue, F., Cheng, Y., Breschi, A., Vierstra, J., Wu, W., Ryba, T., Sandstrom, R., Ma, Z., Davis, C., Pope, B., et al.: A comparative encyclopedia of DNA elements in the mouse genome. Nature 515, 355–364 (2014)

[75] Cunningham, F., Amode, M., Barrell, D., Beal, K., Billis, K., Brent, S., Carvalho-Silva, D., Clapham, P., Coates, G., Fitzgerald, S., et al.: Ensembl 2015. Nucleic Acids Research 43, 662–669 (2015)

[76] Ester, M., et al.: A density-based algorithm for discovering clusters in large spatial databases with noise. In: Kdd, vol. 96, pp. 226–231 (1996)

[77] Gamby, A., et al.: Convex-hull algorithms: Implementation, testing, and experimentation. Algorithms 11(12), 195 (2018)

[78] Pownall, M.E., Miao, L., et al.: Chromatin expansion microscopy reveals nanoscale organization of transcription and chromatin. Science 381(6653), 92–100 (2023)

[79] Chen, K., et al.: Danpos: dynamic analysis of nucleosome position and occupancy by sequencing. Genome Research 23(2), 341–351 (2013)

