## Supplementary Info for "Uncovering High-Resolution Organization of Genomic Loci using Experimentally Informed Polymer Model"

### Addition Information

We develop a simplified multi-scale polymer model to understand the chromatin organization at different resolutions. Our model aims to simulate a 0.2 Mbps long chromatin segment. We use hi-C contact map information at 5kbps resolution to capture global organization. The Hi-C contact map tells about the average contact frequency between different genomic segments among the population of cells. So, we distribute the Hi-C contact information among 100 different conformations of a homo-polymer chain with 40 beads, each representing a 5kbps segment of the target region. Important contacts are distributed among these conformations. A contact is known as an important contact if the contact frequency between any i, j (belonging to 40 beads), is higher than the average frequency($|j-i|$)+ standard deviation of the distribution of frequency($|j-i|$)[1].

$$P_{ij} = \begin{cases} 1, & f_{avg}(|j-i|) \pm f_{std}(|j-i|) < p_{ij} \\ 0, & \text{otherwise.} \end{cases} \tag{1}$$

Here $f_{avg}(|j-i|)$ is the average contact frequency for the genomic segment having the same |j-i| for every i,j having contact probability $p_{ij}$ and $f_{std}(|j-i|)$ stand for standard deviation. The contacts are distributed according to the gaussian probability distribution of contacts among the conformation. For each conformations, we get the contact information to be in contact for each genomic segment bead pair (i,j) as binary matrix. These matrices are used to restrict the dynamics of our simulation for high-resolution copolymer chains representing the target region. The contact information is not enough to generate an ensemble average contact that mimics the Hi-C contact map because dynamics play a major role in manipulating the contact frequency of the simulated ensemble average contact map. Before using this contact information directly we simulate these 100 conformations using a homopolymer chain (Fig. S1) of 40 beads representing the 0.2Mbps segment where each bead represents a 5kbps region. The binary matrix holds the contact information for |j-i| more than 1 as in a polymer chain all the adjacent beads remain always in contact. This helps to know whether the dynamics favour mimicking our global-level organization.

$$U^H_{Total} = U^H_{bond} + U^H_{HICcont} + U^H_{WCA} \tag{2}$$

$$\frac{U^H_{WCA}(r_{ij})}{k_B T} = \begin{cases} 4\left[\left(\frac{d_{ij}}{r_{ij}}\right)^{12} - \left(\frac{d_{ij}}{r_{ij}}\right)^{6}\right] + 1\ ; \ \ r_{ij} < 2^{\frac{1}{6}} d_{ij} \\ \\ 0, \ \text{otherwise.} \end{cases} \tag{3}$$

$$U^H_{HARMO}(r_{ij}) = K_{harmo}(r_{ij} - r_0)^2\ ;\ |j-i| = 1\ ;\ r_0 = 30nm \tag{4}$$

$$U^H_{HICcont}(r_{ij}) = K_{hic}(r_{ij} - r_0)^2\ ;\ |j-i| >= 2\ \&\ P_{ij} = 1\ ;\ r_0 = 30nm \tag{5}$$

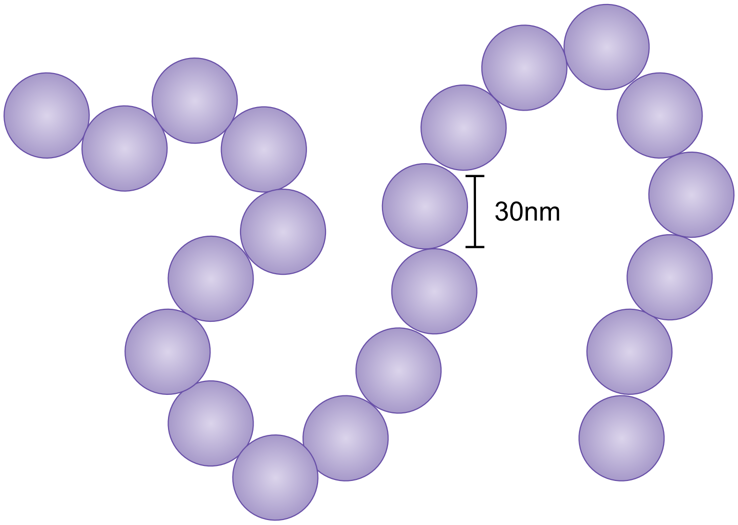


**Figure S1.** Homopolymer chain with each bead diameter 30nm.

In WCA potential $d_{ij}$ is taken 30nm and we put $= K_{harmo}$ bit high so that contacts from binary matrix could not break the chain. We systematically allow contact from a binary matrix to be formed taking care of unwanted chaos. We start from a closer genomic segment (|j-i|) to be in contact with the binary matrix and move in ascending order of |j-i| as the previous pair complete the bond formation. Once all the binary contact is established, we start to dump the structure and calculate the ensemble average contact map. After getting a good agreement with the Hi-C contact map we start our high-resolution simulation with the copolymer chain. The copolymer chain has two types of beads (Fig S2) that differ in size. Beads bigger in size represent the nucleosomes and the beads small in size represent linker DNA. Nucleosome beads represent 142 base pairs and the linker DNA bead represents around 8 base pairs. The position of the nucleosome beads in the copolymer chain is decided using MNase-Seq experimental data using comprehensive bioinformatics pipeline DANPOS[2]. In the experiment, Micrococcal nuclease (MNase) digests non-protein-segment and the fragments left behind are aligned to the DNA sequence, providing a genome-wide position of nucleosome. The polymer chain at almost base pair resolution has a larger number of beads to simulate and cause computational complexity. Further, to overcome this complexity we develop a GPU-based simulator using CudaC programming language with cuda version 11.0 and GeForce GTX10 architecture. This simulator provides 25× speedup than a based simulator to handle around 50,000 beads system. Using the simulator, we simulate our high-resolution copolymer chain of around 10,000 NL beads representing 0.2Mbps chromatin segment. We use Langevin dynamics to simulate our system using a simplified polymer model where every ith bead has position $r_i$ in the polymer chain and $r_{ij}$ is the distance between ith and jth particle. The system has the total energy:

$$U_{Total} = U_{FENE} + U_{WCA} + U_{FENE} + U_{WCA}^{COM} + U_{HICcont}^{COM} \tag{6}$$

The consecutive beads are connected by the finite extensible nonlinear elastic (FENE) potential:

$$U_{FENE}(r_{i,i+1}) = U_{WCA}(r_{i,i+1}) - \frac{K_{FENE}R_0^2}{2} \log\left[1 - \left(\frac{r_{i,i+1}}{R_0}\right)^2\right] \tag{7}$$

Also, consecutive three linker beads are linked using Kratky-Porod potential, which helps them to maintain the persistent length between linker DNA beads. The magnitude of the bending potential manipulates the persistence of linker DNA.

$$U_{BEND}(\theta) = K_{BEND}\left[1 - \cos(\theta)\right] \tag{8}$$

where,

$$\cos(\theta) = (r_i - r_{i-1}) \,.\, (r_{i+1} - r_i) \tag{9}$$

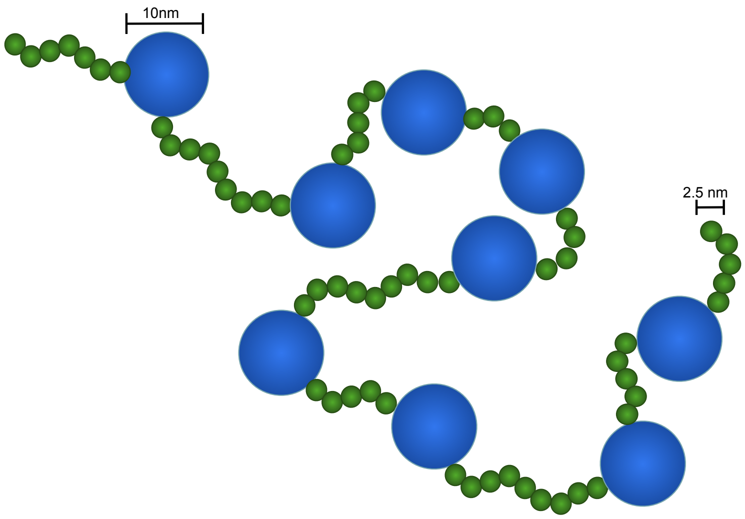


**Figure S2.** Copolymer chain with two types of bead; small size bead have a diameter of 2.5nm and the bead bigger in size have a diameter of 10nm.

Here $\theta$ is the angle formed by three consecutive linker DNA beads and $K_{BEND} = l_p k_B T$ for linker DNA beads. $l_p$ is the persistence length considering 147 linker base pairs to behave like a rod, resulting in a length of around 50nm. We use $\sigma$ to represent the size of the linker DNA bead. In the unit of $\sigma$, $l_p$ is $20\sigma$. To avoid overlapping between non-bounded beads, we use Weeks-Chandler-Andersen(WCA) potential taking care of all the steric interactions.

$$\frac{U_{WCA}(r_{ij})}{k_B T} = \begin{cases} 4\left[\left(\frac{d_{ij}}{r_{ij}}\right)^{12} - \left(\frac{d_{ij}}{r_{ij}}\right)^{6}\right] + 1, & \frac{r_{ij}}{2.5} < 2^{\frac{1}{6}} d_{ij} \; and \; d_{ij} = 10nm \\ 4\left[\left(\frac{d_{ij}}{r_{ij}}\right)^{12} - \left(\frac{d_{ij}}{r_{ij}}\right)^{6}\right] + 1, & r_{ij} < 2^{\frac{1}{6}} d_{ij} \; and \; d_{ij} \neq 10nm \\ 0, & \text{otherwise.} \end{cases} \tag{10}$$

In our copolymer chain, the nucleosomal bead has a diameter of 10nm and linker DNA bead has a diameter of 2.5nm[3] as each base pair has a size of 0.34. Here $d_{i,j}$ refers to the average diameter of interacting beads and $r_{i,j}$ is the distance between $r_i$ and $r_j$. In eq. (7) is $R_0$ is $1.6 d_{i,j}$. To implement the binary contact matrix in our high-resolution copolymer chain, we take care of a list which tell about NL beads index related to 5kbps index from homopolymer chain in our target region. With the help of center of mass position (COMP) of NL beads for each 5kbps region we maintain the reference distance between every COMP. If two 5kbps are in contact they start to maintain reference distance between COMP pair. In case of they have $P_{ij} = 0$ they follow excluded volume potential to avoid segmental overlapping. Each 5kbps region have a spherical space of diameter 90nm following nucleus base pair density to behave NL beads purely entropically.

$$\frac{U^{COM}_{WCA}(r^{COM}_{ij})}{k_B T} = \begin{cases} 4\left[\left(\frac{r^{COM}_0}{r^{COM}_{ij}}\right)^{12} - \left(\frac{r^{COM}_0}{r^{COM}_{ij}}\right)^{6}\right] + 1 \; ; \; r^{COM}_{ij} < 2^{\frac{1}{6}} r^{COM}_0 \\ 0, \; \text{otherwise.} \end{cases} \tag{11}$$

$$U^{COM}_{HICcont}(r^{COM}_{ij}) = K^{COM}_{hic}(r^{COM}_{ij} - r^{COM}_0)^2 \; ; \; |j-i| >= 2 \; \& \; P_{ij} = 1 \; ; \; r^{COM}_0 = 90nm \tag{12}$$

During the simulation, the position of beads is updated by following the Langevin equation

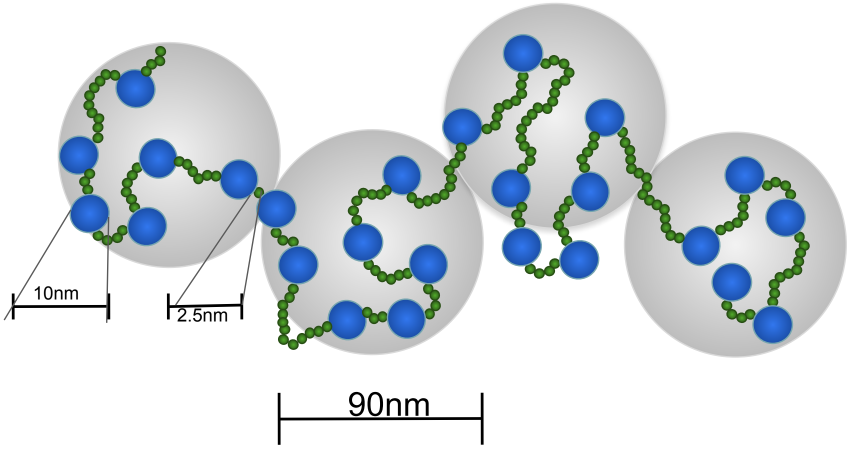


**Figure S3.** Each 5kbps segment of copolymer chain of NL beads maintains a center of mass of 90nm. The NL beads almost mimic a spherical volume of 90nm diameter.

$$m_i \frac{d^2\mathbf{r}_i}{dt^2} = -\nabla U_i - \gamma_i \frac{d\mathbf{r}_i}{dt} + \sqrt{2k_B T \gamma_i}\eta_i(t) \tag{13}$$

Here $m_i$ is mass of respective bead, $r_i$ is the position vector, $\gamma_i$ is the fraction felt due to implicit solvent and $\eta_i(t)$ is uncorrelated random noise that follow the equation mentioned below

$$\langle \eta_\alpha(t) \rangle = 0 \tag{14}$$

$$\langle \eta_\alpha(t)\eta_\beta(t^{'}) \rangle = \delta_{\alpha\beta}\delta(t-t^{'}) \tag{15}$$

The noise is controlled by thermal fluctuation in the form of thermal energy as a multiplication of Boltzmann coefficient ($k_B$) and temperature (T). Here with the reduced unit parameter, we use T=1. The $U_i$ is the summation of all the potential representing the polymer chain interaction. Here we consider the mass and fraction coefficient for all the beads identical. To solve the above equation we use the velocity-Verlet algorithm with the time step $\Delta t = 0.005\tau$ where $\tau$ is the simulation time unit. We initialize the simulation with a linear chain. We provide inter nucleosome potential using eq.(11) and the contact information we distributed in 100 conformation from Hi-C contact map put constrain among NL beads to follow the connection between 5kbps regions. We simulate for 50000 $\tau$ and after 17500 $\tau$ we start to dump the structure for every 250 $\tau$. Our system satisfies the restriction imposed to capture global organization details within initial 17500 $\tau$. We run 100 simulation with respect to global organization information distributed among 100 conformation. Each simulation provide 130 conformation and we get 13000 structures after simulating the system for all the restriction layout.

### Experimental data for modelling and validation

We export the Hi-C contactmap[4] using 4DNUCLEOME database. We use experimental set 4DNES2M5JIGV with sample H1-hESC (Tier 1) from experimental type in situ Hi-C. Similar,[4] from experiment set 4DNES21D8SP8 we get micro-C contact map for the same sample for experimental type. We get the nucleosome position data for our target system from nucmap database that provide genome-wide position map of nucleosomes for different species. For H1 cell line with experiment component GSM1194220 for sample id hsNuc0070101 we get the position of nucleosomes for our target region.

### ChromHMM

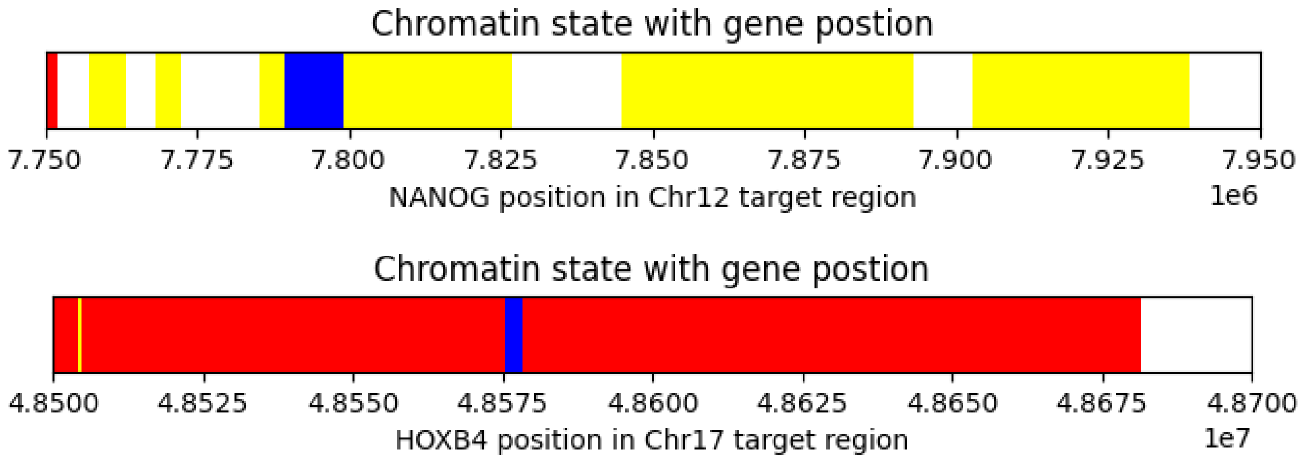


**Figure S4.** ChromHMM differentiate between the transcription active region and inactive region. The red colour shows the transcriptionally inactive region, the yellow colour shows the transcriptionally active region, the blue colour shows the position of the gene position and the uncoloured space is quiescent locations that could not be defined based on transcription activity.

We aim to understand the chromatin organization at different resolutions for a eukaryotic chromatin region. Initially, we simulate for HoxA1 gene loci chromatin region 0.2Mbps region. After successfully capturing the chromatin organization detail to understand the organization difference based on transcription activity, we chose two more different gene loci systems Nanog and HoxB4. We use chromHMM[5] tool to differentiate between their transcription activity which suggests that Nanog is transcriptionally active and HoxB4 is a transcriptionally inactive gene loci(Fig. S4).

### Contact map and boundary prediction

We simulate three target systems with gene loci HoxA1, Nanog and HoxB4. As a simulation result, we get 13,000 structures for each system. We calculate the ensemble average contact map using these structures. To capture the contact map we coarse-grain the high-resolution copolymer chain into a desired resolution homopolymer chain. To calculate a contact up for the respective resolution we randomly select a genomic pair to be in contact for a random structure and if a random number is smaller than $exp(-\frac{r_{ij}^2}{r_{cut}^2})$ then pair is considered to be in contact. up all the The contact map gives the interaction frequency for each pair of genomic regions and forms domain-like structures along the diagonal of the map. To predict the boundaries of these domains we use slide box algorithm[3,6]. In this method, we sum interaction frequencies of unique pairs in a range and calculate the contact score for each position.

$$score_k = \frac{1}{2d} \sum_{i=k-d+1}^{k} \sum_{j=k+1}^{k+d} x_{ij} \qquad for \quad d < k < N - d \tag{16}$$

After getting a score, we compare the score of the specific position k to $k+K$ upstream and $k+K$ downstream positions scores. If the the target position has the lowest score then it is caller a boundary. The value of k,d and K are chosen based on contact map visualisation that fit better for the contact map domains.

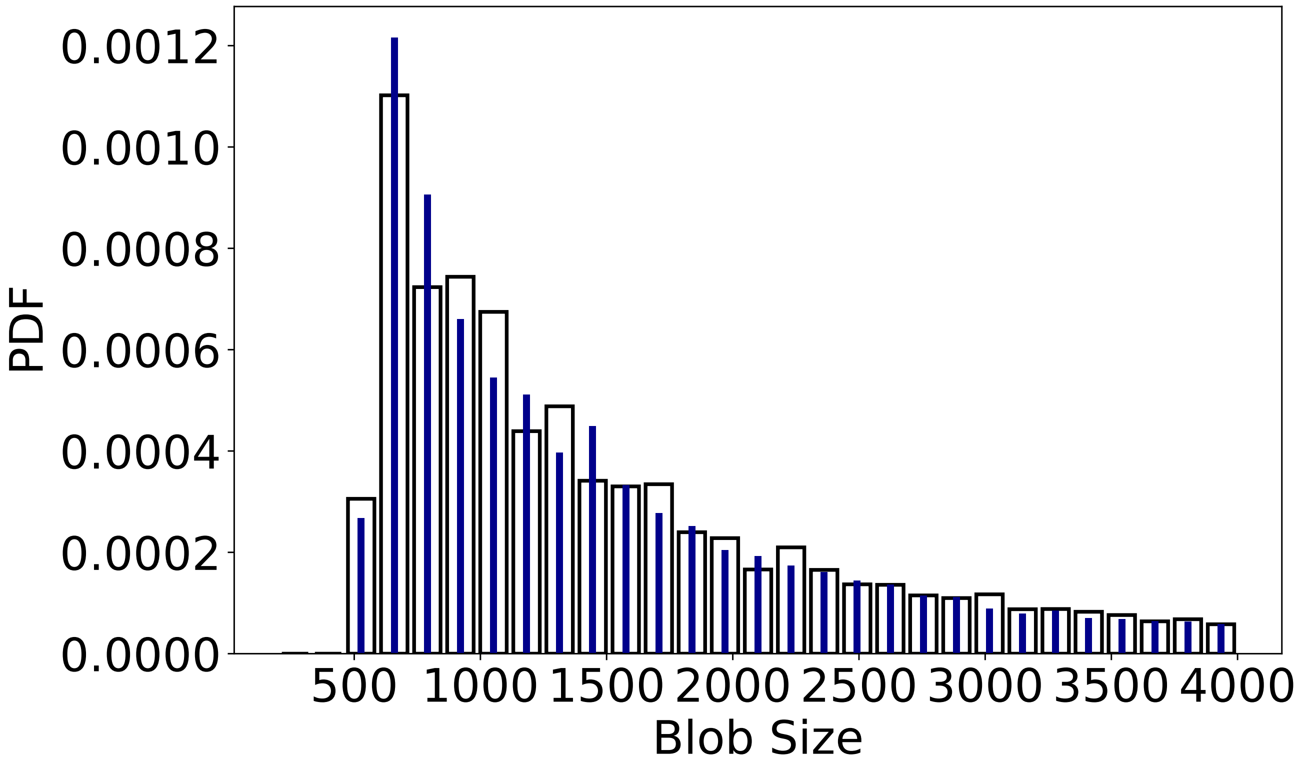


**Figure S5.** Histogram of different blob sizes.

### Blob dyanmics

To understand the behavior of the blobs, we further analyze the conformations obtained during the simulation. We calculate the Nearest Neighbor Distance (NND) of the blobs using the kth neighbouring distance (k = 1 to 40). The observed pattern closely resembles that reported by Barth et al.[7].

To further explore the blobs's dynamics, we compute the flow magnitude of blobs, which quantifies their movement between consecutive time steps. Additionally, we examine the variation in distances between different-sized blobs by calculating the radial distribution of surface area. We then analyze the spatial-temporal auto- and cross-correlation between surface area, NND, and flow magnitude to understand their interdependencies.

The autocorrelation of surface area indicates a high correlation at time lag zero, which quickly diminishes by time lag 4 and continues to decrease over a 40-unit time lag. This suggests that the surface area of blobs has minimal memory of its previous states. Spatially, the effect of surface area diminishes beyond 250 nm, whereas NND and flow magnitude exhibit some dependence on spatial lag, though the effect remains weak. The temporal analysis reveals that both NND and flow magnitude retain memory from previous time steps, persisting longer than surface area.

The spatial-temporal autocorrelation of flow magnitude closely mirrors that of NND, and their cross-correlation further supports this similarity, indicating that blob movement is more pronounced in regions where NND is higher. This suggests that spatially dispersed blobs exhibit greater movement than those that are more compact. Furthermore, the spatial-temporal cross-correlation between surface area and flow magnitude reveals that at a 250 nm spatial lag, blob motion has the greatest impact on blob size. This implies that the dynamics of blobs separated by approximately 250 nm influence surface area the most, with some dependency on previous time steps. However, as the distance decreases or exceeds 250 nm, this effect diminishes, along with its temporal memory.

Finally, the autocorrelation of surface area suggests that up to a 250 nm spatial lag, the surface area of one blob can influence that of others. However, beyond this range, the effect decreases, while flow magnitude—representing blob dynamics—plays a dominant role in determining blob size. Notably, at 250 nm, the influence of flow magnitude is most pronounced, but it weakens as spatial lag increases.

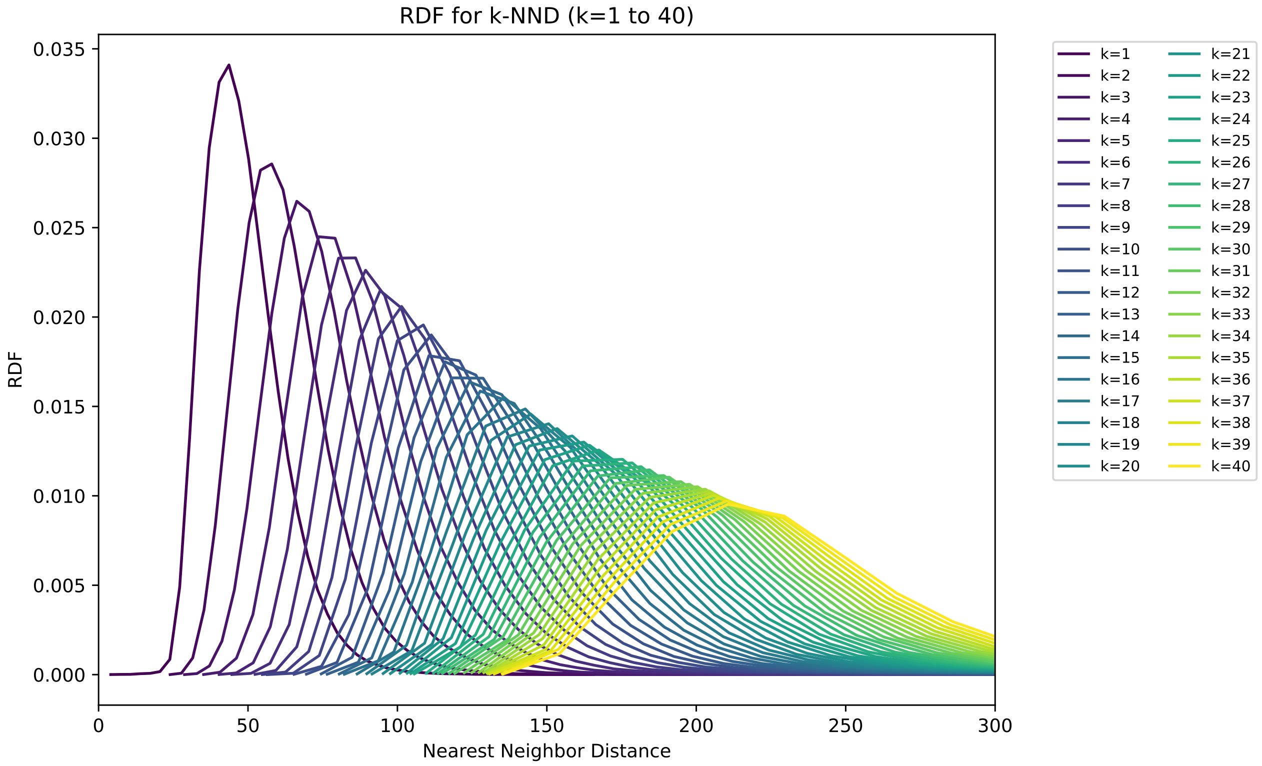


**Figure S6.** Radial distribution of kth nearest neighbor distance.

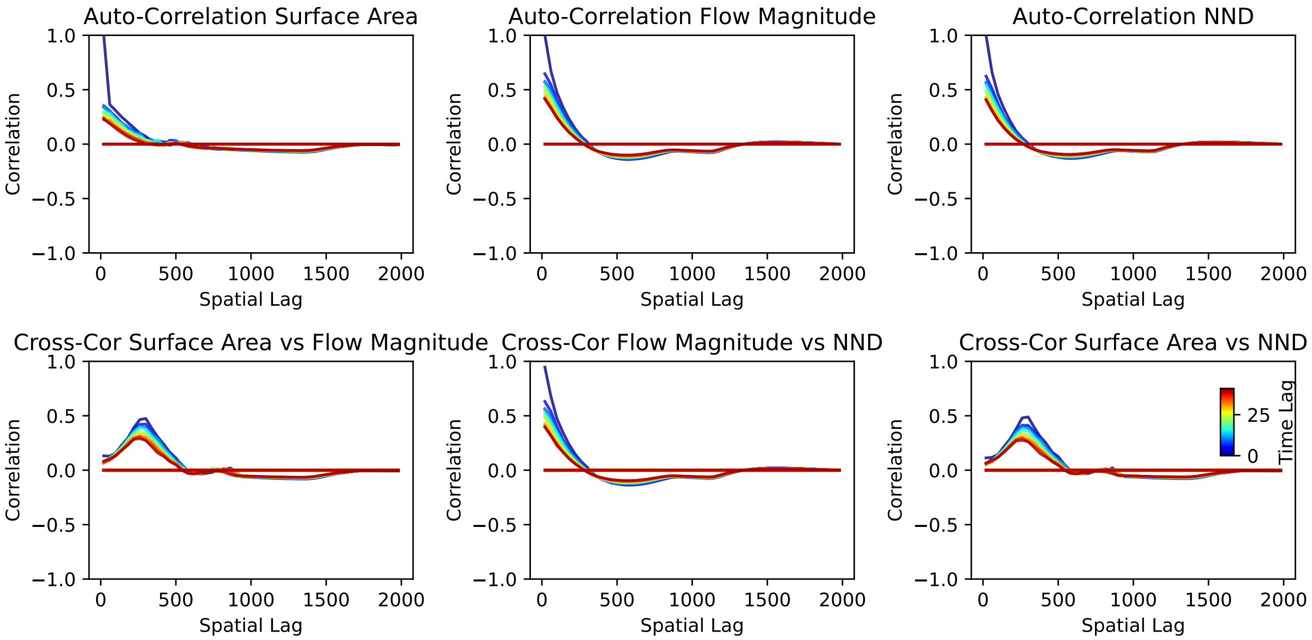


**Figure S7.** Spatial-temporal auto and cross correlation between flow magnitude, NND and surface area of blobs.

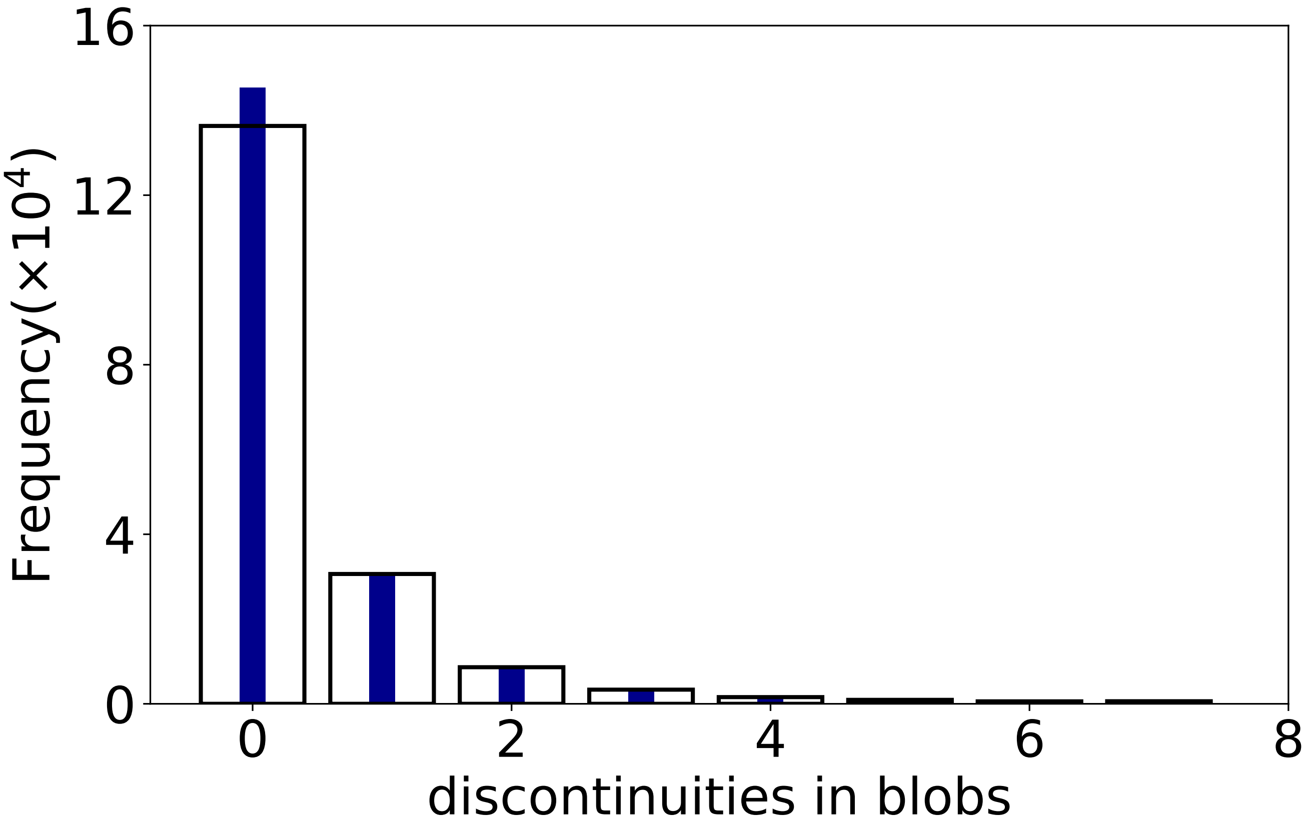


**Figure S8.** Nucleosomes come together and form a blob. A sequential genomic segment forming a blob includes the sequential nucleosomes. But when distant genomic loci come closer and form a blob it can include N multiple sequential nucleosomes. These N multiple sequential nucleosomes suggest an N-1 break between sequential nucleosomes. The plot gives the histogram of breaks in the blobs and suggests that the blobs favour fewer breaks. The blue colour shows the histogram for the Nanog and the black border line shows the histogram for HoxB4.

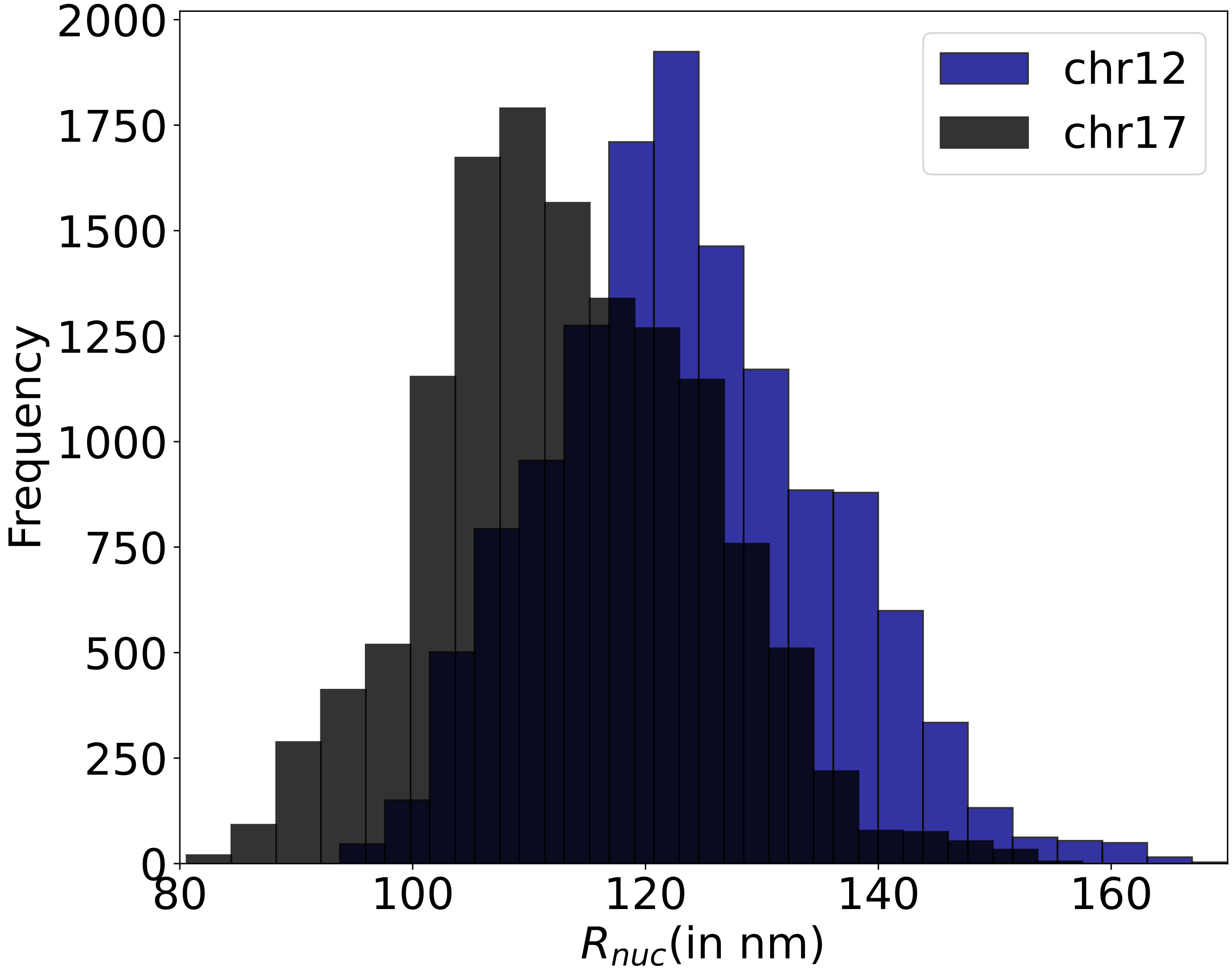


**Figure S9.** The plot shows the histogram of the mean distance between the nucleosomes of the 100 nucleosomes around the gene position in the target region to understand the local level compaction around the gene position. The blue colour shows the histogram for the Nanog gene loci and the black colour shows the histogram for the HoxB4 gene loci and the dark black in the plot shows the overlapping region of the histogram formed by both the Nanog and HoxB4 genes. The plot suggests that Nanog has more spacing than HoxB4 with more distance between the nucleosomes.

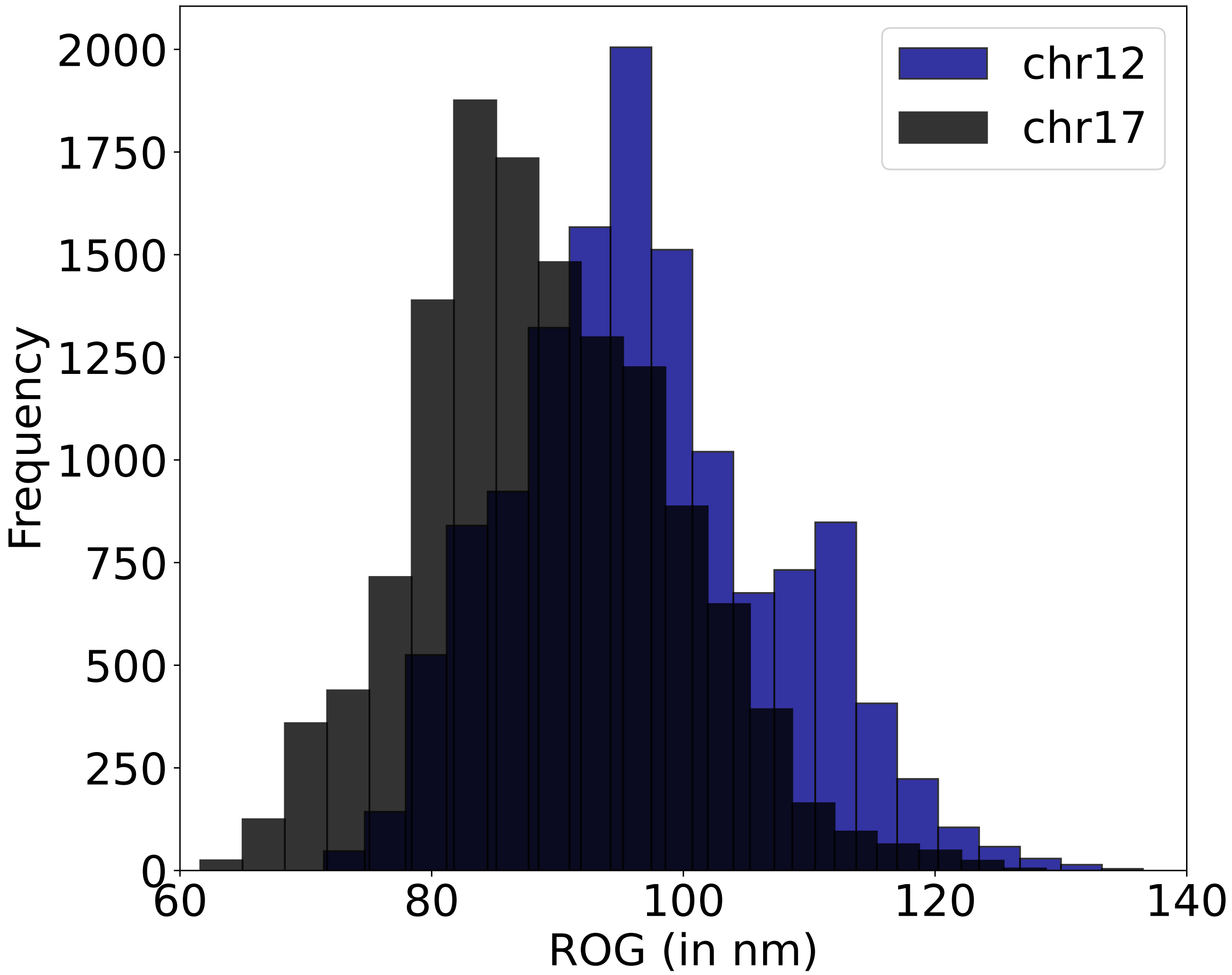


**Figure S10.** The plot shows the histogram for the radius of the gyration of the 100 nucleosomes around the gene position in the target region to understand the local level compaction around the gene position. The blue colour shows the histogram for the Nanog gene loci and the black colour shows the histogram for the HoxB4 gene loci and the dark black in the plot shows the overlapping region of the histogram formed by both the Nanog and HoxB4 genes. The plot shows that the Nanog has a larger ROG than HoxB4 suggesting nucleosomes around gene position in Nanog are distributed in a larger space than HoxB4.

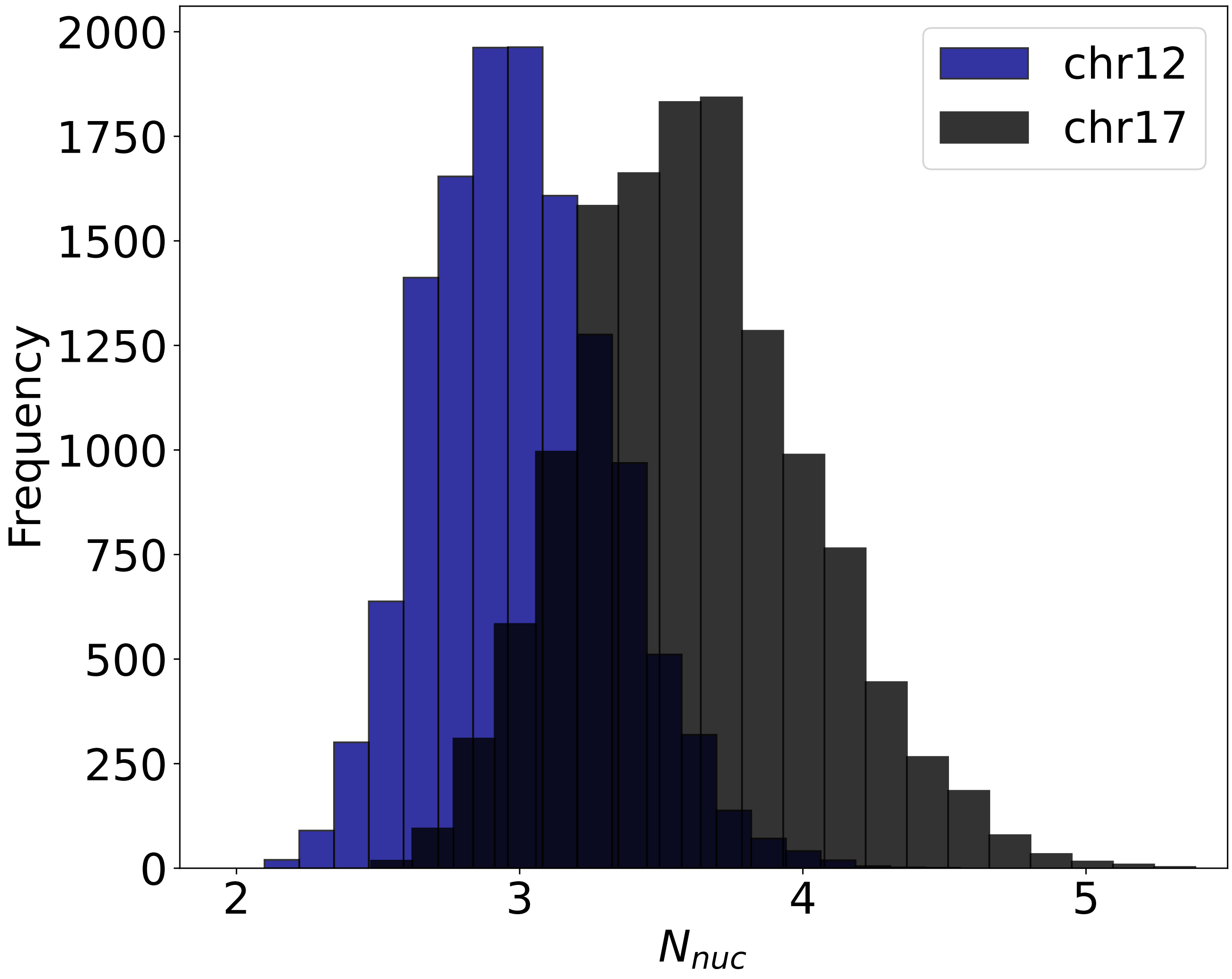


**Figure S11.** The plot shows the histogram for neighbouring nucleosomes with a cutoff of 24nm around every nucleosome in the set of 100 nucleosomes around the gene position in the target region to understand the local level compaction around the gene position. The blue colour shows the histogram for the Nanog gene loci and the black colour shows the histogram for the HoxB4 gene loci and the dark black in the plot shows the overlapping region of the histogram formed by both the Nanog and HoxB4 genes. The plot shows neighbouring nucleosomes in Nanog are lesser than HoxB4 suggesting clusters of nucleosomes around gene position are smaller in Nanog than HoxB4.

### References


1. et al., K. S. Predicting scale-dependent chromatin polymer properties from systematic coarse-graining. *Nat Commun* **14**, 4108 (2023).
2. Chen, K. & others. Danpos: dynamic analysis of nucleosome position and occupancy by sequencing. *Genome Res.* **23**, 341–351 (2013).
3. Wiese, O. e. a. Nucleosome positions alone can be used to predict domains in yeast chromosomes. *Proc. Natl. Acad. Sci. USA* **116**, 17307–17315 (2019).
4. et al., K. N. Ultrastructural details of mammalian chromosome architecture. *Mol Cell* **78**, 554–565.e7 (2020).
5. et al., E. J. Chromhmm: automating chromatin-state discovery and characterization. *Nat Methods* **9**, 215–216 (2012).
6. Hsieh, T. *et al.* Mapping Nucleosome Resolution Chromosome Folding in Yeast by Micro-C. *Cell* **162**, 108–119 (2015).
7. Barth, R. *et al.* Coupling chromatin structure and dynamics by live super-resolution imaging. *Sci. Adv.* **6**, eaaz2196 (2020).
